# Comparative Genome Analysis of *Scutellaria baicalensis* and *Scutellaria barbata* Reveals the Evolution of Active Flavonoid Biosynthesis

**DOI:** 10.1101/2020.02.18.954164

**Authors:** Zhichao Xu, Ranran Gao, Xiangdong Pu, Rong Xu, Jiyong Wang, Sihao Zheng, Yan Zeng, Jun Chen, Chunnian He, Jingyuan Song

## Abstract

*Scutellaria baicalensis* and *Scutellaria barbata*, common medicinal plants of the Lamiaceae family, produce specific flavonoid compounds with antioxidant and antitumor activities, including baicalein, scutellarein, norwogonin, wogonin, and their glycosides. Here, we reported two chromosome-level genome assemblies of *S. baicalensis* and *S. barbata* with significant quantitative chromosomal variation (2n = 18 and 2n = 26, respectively). The divergence of *S. baicalensis* and *S. barbata* occurred far earlier than previously reported, and a whole-genome duplication event was identified. The insertion of long terminal repeat elements after speciation might be responsible for the observed chromosomal expansion and rearrangement. The comparative genome analysis of congeneric species elucidated the species-specific evolution of chrysin and apigenin biosynthetic genes, such as the *S. baicalensis*-specific tandem duplication of the phenylalanine ammonia lyase (PAL) and chalcone synthase (CHS) genes, and the *S. barbata*-specific duplication of 4-CoA ligase (4CL) genes. In addition, the paralogous duplication, collinearity, and expression diversity of CYP82D subfamily members revealed the functional divergence of flavone hydroxylase genes between *S. baicalensis* and *S. barbata*. These *Scutellaria* genomes highlight the common and species-specific evolution of flavone biosynthetic genes, promoting the development of molecular breeding and the study of the biosynthesis and regulation of bioactive compounds.

## Introduction

Plant-specific flavonoids, including flavones, flavonols, anthocyanins, proanthocyanidins, and isoflavones, play important functions in plants, such as flower pigmentation, UV protection, and symbiotic nitrogen fixation [1−3]. Flavonoid metabolites also have biological and pharmacological activities in human health, including antibacterial and antioxidant functions, and the treatment of cancer, inflammatory, and cardiovascular diseases [3]. The genus *Scutellaria*, belong to the Lamiaceae family, comprises common herbal plants enriched by bioactive flavonoids, and approximately 300 to 360 *Scutellaria* species have been documented as having the characteristic flower form of upper and lower lips [4,5]. Only two *Scutellaria* species, *Scutellaria baicalensis* and *Scutellaria barbata*, are recorded in the Chinese pharmacopoeia, and the roots of *S. baicalensis* and dried herbs of *S. barbata* are the basis of the Chinese medicines *Huang Qin* and *Ban Zhi Lian*, respectively, which have been well known heat-clearing and detoxifying herbs for thousands of years [6]. The main biologically active compounds in *Scutellaria* are derivatives of chrysin and apigenin, such as baicalein, scutellarein, wogonin, and their glycosides (baicalin, scutellarin, and wogonoside) [7−10]. Baicalin has been confirmed to activate carnitine palmitoyltransferase 1 in the treatment of diet-induced obesity and hepatic steatosis, leading to extensive interest in the potential antilipemic effect of this compound [11,12].

Illuminating the chemodiversity and biosynthesis of the active constituents of *Scutellaria* will provide a foundation for investigating the use of *Huang Qin* and *Ban Zhi Lian* in traditional Chinese medicine (TCM), and the production of these natural products via synthetic biology [13]. In *S. baicalensis*, the biosynthetic genes of the root-specific compounds baicalein and norwogonin have been functionally identified, providing an important basis for studying the biosynthesis and regulation of the natural products that make up *Huang Qin* [14,15]. Recently, the *in vitro* production of baicalein and scutellarein in *Escherichia coli* and *Saccharomyces cerevisiae* has been carried out based on the guidance of synthetic biology [16,17], but the metabolic engineering of these compounds still faces considerable challenges, including the discovery and optimization of biological components. The *Salvia miltiorrhiza* genome from the Lamiaceae family provides useful information associated with secondary metabolism for the rapid functional identification of biosynthetic and regulatory genes [18−23]. In contrast, the genomes of the *Scutellara* genus remains unclear, and the reliance on transcriptome data from short-read sequencing has restricted gene discovery and analyses of genome evolution, including studies of gene family expansion and contraction, the evolution of biosynthetic genes, and identification of regulatory elements [24].

Significant morphological differences are present at the macroscopic level between *S. baicalensis* and *S. barbata*; these species are differentiation is mainly characterized by the fleshy rhizome and branched stem of *S. baicalensis* and the fibrous root and erect stem of *S. barbata*. The active compounds baicalein, wogonin and scutellarein are differentially distributed in the roots and aerial parts of *S. baicalensis* and *S. barbata*. Here, we performed *de novo* sequencing and assembly of the *S. baicalensis* and *S. barbata* genomes using a long-read strategy and Hi-C technology. The chromosome-level genome of *S. baicalensis* and *S. barbata* revealed their divergence time, chromosomal rearrangement and expansion, whole-genome duplication, and the evolutionary diversity of flavonoid biosynthesis. The study provided significant insights for the molecular assisted breeding of important TCM resources, genome editing, and understanding the molecular mechanisms of the chemodiversity of active compounds.

## Results and discussion

### High-quality genome assemblies and annotation

The size of the *S. baicalensis* genome was predicted to be 440.2 ± 10 Mb and 441.9 Mb using flow cytometry and the 21 *k*-mer distribution analysis (approximately 0.96% heterozygosity) (**Figure 1**A, Figure S1). The genome survey of *S. barbata* showed a 404.6 Mb genome size and 0.28% heterozygosity via the 21 *k*-mer distribution analysis (Figure 1A, Figure S1). Third-generation sequencing platforms, including PacBio and Oxford Nanopore technologies, have been confirmed to have a significant advantage in *de novo* assembly and in processing data with complex structural variation due to high heterozygosity and repeat content [25−27]. Thus, 52.1 Gb Oxford Nanopore technology (ONT) reads (∼120 ×) with an N50 of 16.3 kb from *S. baicalensis* and 51.7 Gb single molecule, real-time sequencing (SMRT) reads from the PacBio platform (∼130 ×) with an N50 of 9.8 kb from *S. barbata* were produced to assemble highly contiguous genomes (Table S1). The low-quality long reads were further corrected and trimmed to yield 20.2 Gb ONT reads with an N50 of 35.5 kb from *S. baicalensis* and 18.0 Gb SMRT reads with an N50 of 15.3 kb from *S. barbata* using the CANU pipeline.

**Figure 1.**
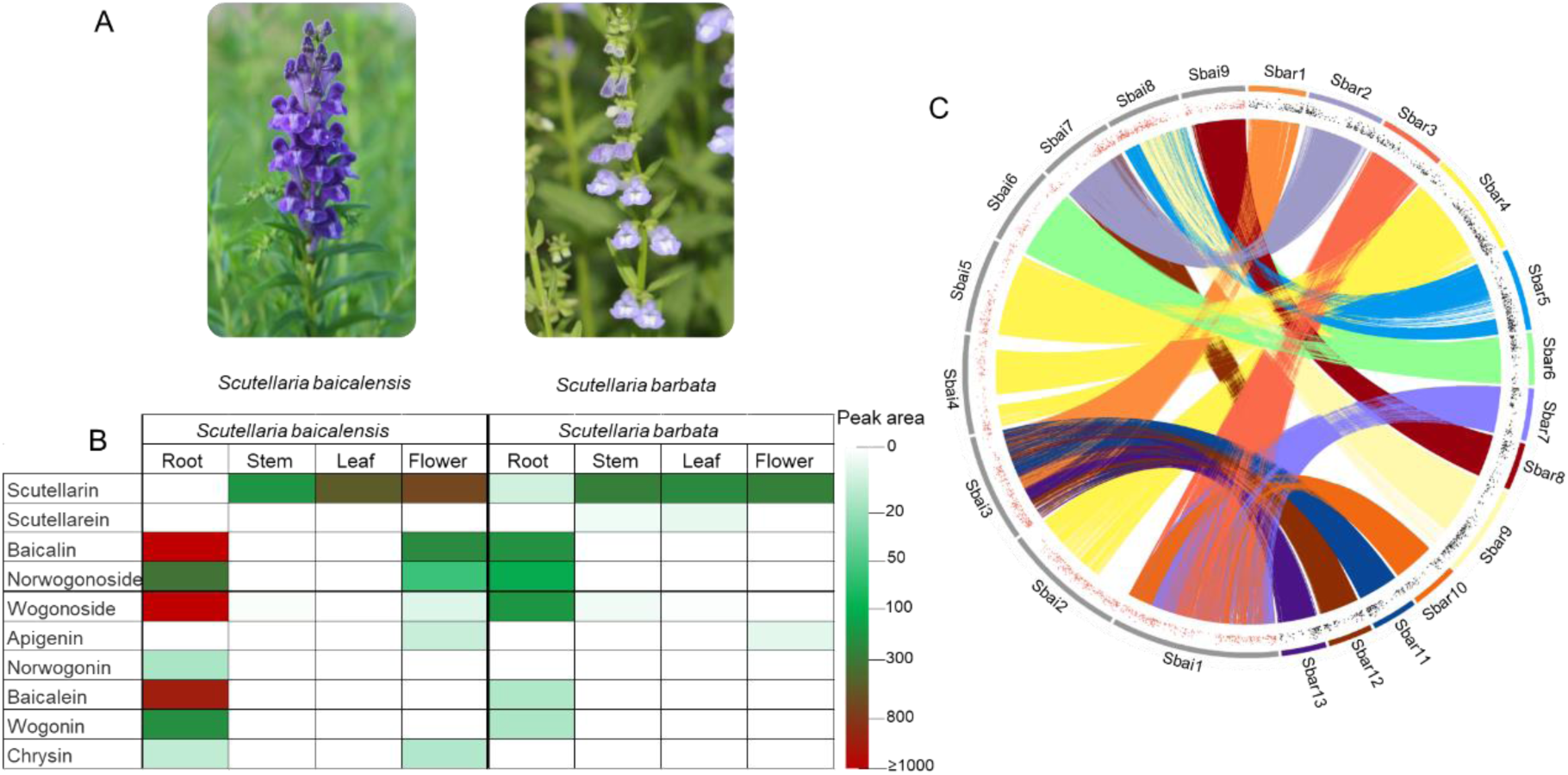
The similar morphology and flavonoid distribution of *S. baicalensis* and *S. barbata*, as well as their genome collinearity. A. Morphological differences between the flowers of *S. baicalensis* and *S. barbata*. B. Content distribution of flavone compounds in different tissues of *S. baicalensis* and *S. barbata*, including roots, stems, leaves and flowers. C. Comparison of nucleic acid sequences from 9 *S. baicalensis* chromosomes and 13 *S. barbata* chromosomes; mapping regions with more than 90% sequence similarity over 5 kb were linked. The red and black dots represent significant changes in gene expression (Log_2_foldchange>1, FPKM>10) in the root tissues of *S. baicalensis* and *S. barbata*, respectively.

The contiguous assembly of the *S. baicalensis* and *S. barbata* genomes was performed using the optimized SMARTdenovo and 3 × Pilon polishing (50 × Illumina reads) packages. For *S. baicalensis*, the contig-level genome assembly, which was 377.0 Mb in length with an N50 of 2.1 Mb and a maximum contig length of 9.7 Mb covered 85.3% of the estimated genome size (Table S2). The *S. baicalensis* genome identified 91.5% of the complete Benchmarking Universal Single-Copy Orthologs (BUSCO) gene models and had an 88.7% DNA mapping rate, suggesting a high-quality genome assembly. For *S. barbata*, the contiguous contig assembly of 353.0 Mb with an N50 of 2.5 Mb and maximum contig of 10.5 Mb covered 87.2% of the predicted genome size (Table S2). The *S. barbata* genome identified 93.0% of complete BUSCO gene models and had a 95.0% DNA mapping rate. The high-quality genome assemblies of *S. baicalensis* and *S. barbata* showed the great advantage of single molecule sequencing, with assembly metrics that were far better than those of other reported genomes of Lamiaceae species, i.e., *Salvia miltiorrhiza* [28] and *Mentha longifolia* [29].

Given the assembly continuity, with a contig N50 of over 2 Mb for the *S. baicalensis* and *S. barbata* genomes, chromosome conformation capture (Hi-C) technology was applied to construct chromosome-level genomes [30]. In total, 99.8% and 98.8% of the assembled *S. baicalensis* and *S. barbata* contigs were corrected and anchored to 9 and 13 pseudochromosomes (2n = 18 for *S. baicalensis*, 2n = 26 for *S. barbata*) using a Hi-C interaction matrix with N50 values of 40.8 Mb and 23.7 Mb, respectively. The strong signal along the diagonal of interactions between proximal regions suggested that the Hi-C assemblies for the *S. baicalensis* and *S. barbata* genomes had high quality (Figure S2).

The *S. baicalensis* genome comprised 33,414 protein-coding genes and 2,833 noncoding RNAs (ncRNA), and 41,697 genes and 1,768 ncRNAs were annotated in the *S. barbata* genome (Table S4). Consistent with the genome assembly quality assessment, orthologs of 93.2% and 94.3% of the eukaryotic BUSCOs were identified in the *S. baicalensis* and *S. barbata* gene sets, suggesting the completeness of the genome annotation (Table S4). The gene-based synteny between *S. baicalensis* and *S. barbata* showed chromosome number variation and structural rearrangement (Figure 1C, Figure S3, Table S3). In addition, the alignment at the DNA sequence level also showed the large-scale structural variations between *S. baicalensis* and *S. barbata* genome (Figure S4).

### Chromosome rearrangements and expansion after speciation

Transposable elements (TEs) accounted for approximately 55.2% (208,004,279) and 53.5% (188,790,851) of the *S. baicalensis* and *S. barbata* genomes, respectively (Table S5 and S6). And, 57.6% and 59.9% of these TEs were long terminal repeat (LTR) elements, respectively. Furthermore, we identified 1,225 and 1,654 full-length LTR elements, including *Gypsy* (342 and 310) and *Copia* (354 and 618) elements, in the *S. baicalensis* and *S. barbata* genomes (Table S7). However, there were significant differences in the insertion times of LTR elements, indicating that the LTRs (1.41 MYA, million years ago) in *S. baicalensis* are more ancient than those (0.88 MYA) in *S. barbata*, assuming a mutation rate of μ=1.3×10^−8^ (per bp per year) (Figure S5, Table S7). The recent insertion and activation of LTRs might be key factors in the generation of chromosome rearrangements and expansion of *S. barbata* [31,32]. The ribosomal RNAs (rRNAs) and simple sequence repeats (SSRs) were further annotated (Table S8 and S9). A total of 142,951 and 147,705 SSRs were annotated in *S. baicalensis* and *S. barbata*, respectively, and these SSRs will provide useful molecular markers for breeding and genetic diversity studies.

We employed a genome-wide high-resolution Hi-C interaction analysis of *S. baicalensis* and *S. barbata* to characterize the architectural features of folded eukaryotic chromatin, including interchromosomal interactions, the compendium of chromosomal territories, and A/B compartments [33−35]. First, 159 × and 173 × Hi-C sequencing reads were uniquely mapped (49.6% and 59.0%) to the *S. baicalensis* and *S. barbata* reference genomes, respectively. Then, 84.8 and 113.1 million valid interaction pairs were obtained to construct the matrix of interactions among 100 kb binned genomic regions across all 9 *S. baicalensis* chromosomes and 13 *S. barbata* chromosomes. The whole-chromosome interactions of *S. baicalensis* indicated that chr5 and chr9 had a closer association than the other chromosome pairs. In *S. baicalensis*, the chromosome set including chr2, chr3 and chr8 showed enrichment and association with each other, and depletion with other interchromosomal sets, implying that these three chromosomes were mutually closer in space than the other chromosomes (Figure S6). In *S. barbata*, the chromosomal territories of chr4, chr5, and chr9, with mutual interactions, occupied an independent region in the nucleus (Figure S7).

As the secondary major structural unit of chromatin packing in *S. baicalensis* and *S. barbata*, the A/B compartments representing open and closed chromatin, respectively, were characterized according to an eigenvector analysis of the genome contact matrix. Similarly, more than half of the assembled *S. baicalensis* and *S. barbata* genomes (53.2% and 52.0%) were identified as A compartment in the leaf tissue. As expected, the TE density in the A compartment was dramatically lower than that in the B compartment (*p* < 0.001), and the gene number per 100 kb was significantly higher in the A compartment (*p* < 0.001) (Figure S5 and S6), indicating a positive correlation between the A compartment and transcriptional activity or other functional measures [33,35].

### Whole-genome duplication events between *S. baicalensis* and *S. barbata*

Conserved sequences, including orthologs and paralogs, can be used to deduce evolutionary history based on whole-genome comparisons. Here, orthologous groups of amino acid sequences from 11 angiosperms were identified, yielding a total of 19,479 orthologous groups that covered 291,192 genes. Among these, 120,459 genes clustering into 6,837 groups were conserved in all examined plants. Computational analysis of gene family evolution (CAFÉ) showed that 1,180 and 1,853 gene families were expanded in the *S. baicalensis* and *S. barbata* lineages, respectively, while 1,599 and 1,632 gene families contracted, respectively (Figure 2A, Figure S8, Table S10). Functional exploration of *Scutellaria*-specific genes indicated that domains related to secondary metabolite biosynthesis, such as transcription factors, cytochrome P450s, and O-methyltransferase were markedly enriched.

**Figure 2.**
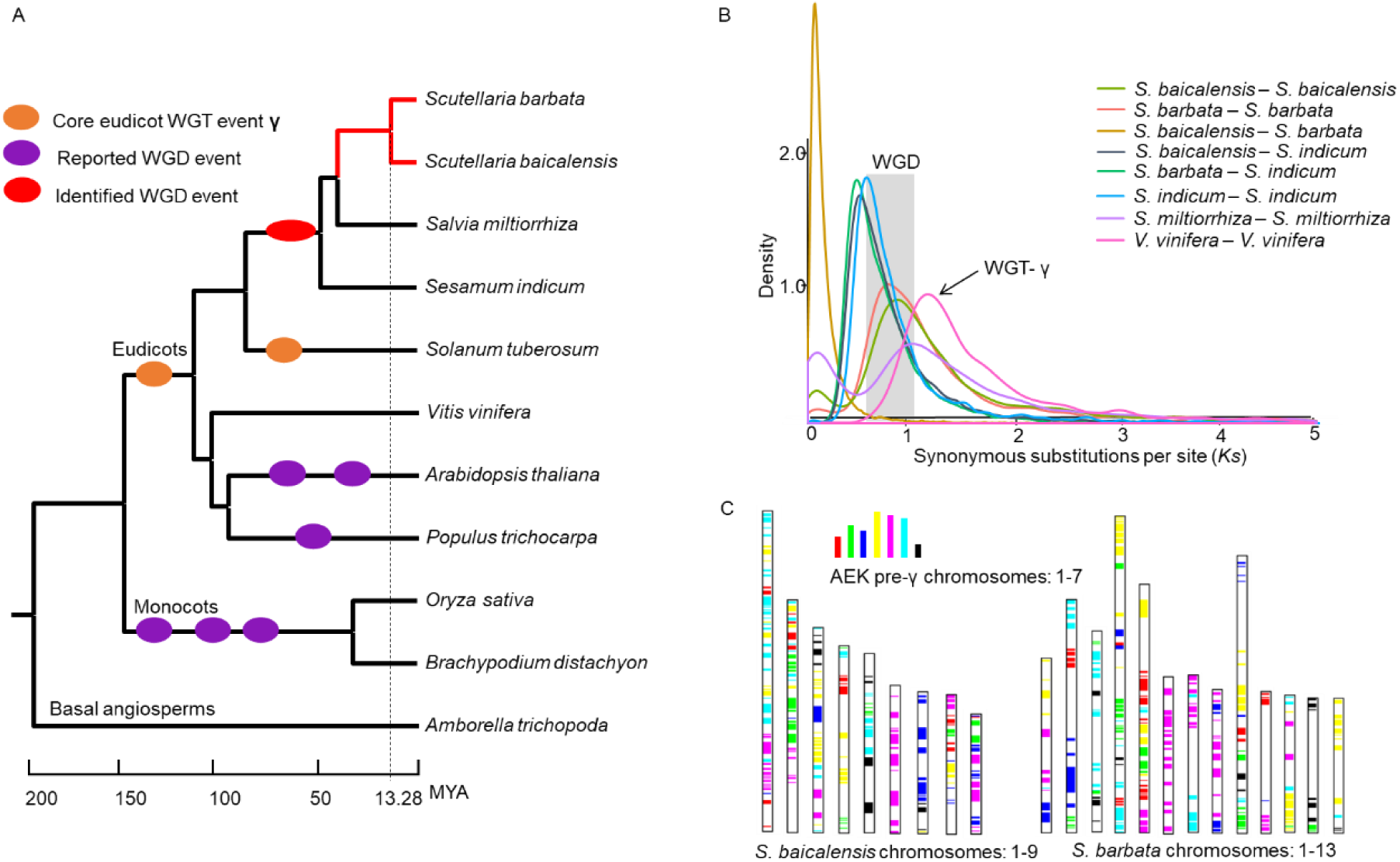
The divergence time and whole genome duplication of the *S. baicalensis* and *S. barbata* genomes. A. The phylogenetic tree was constructed using 235 single-copy orthologous genes from 11 angiosperms. The basal angiosperm *Amborella trichopoda* was chosen as the outgroup. Speciation times were estimated based on the reported divergence times *Amborella_trichopoda*-*Vitis_vinifera* (173-199 MYA) and *Populus trichocarpa*-*Arabidopsis thaliana* (98-117 MYA). The orange ovals represented the reported whole genome triplication events (WGT), and the red and purple ovals represent whole genome duplication events (WGD). B. Synonymous substitution rate (*Ks*) distributions of syntenic blocks for the paralogs and orthologs of *S. baicalensis, S. barbata, S. miltiorrhiza, S. indicum*, and *Vitis vinifera.* C. Comparison with ancestral eudicot karyotype (AEK) chromosomes. The syntenic AEK blocks are painted onto *S. baicalensis* and *S. barbata* chromosomes, respectively.

In addition, 235 single-copy genes in all tested plants were identified and used to construct a phylogenetic tree, indicating that these two *Scutellaria* species were most closely related to *Salvia miltiorrhiza* with an estimated divergence time of 41.01 MYA; *S. baicalensis* and *S. barbata* were grouped into one branch, with an estimated divergence time of approximately 13.28 MYA (Figure 2A). The Phylogenetic tree also supported the close relationship between Lamiaceae (*S. baicalensis, S. barbata* and *S. miltiorrhiza*) and Pedaliaceae (*Sesamum indicum*) with the divergence time of approximately 49.90 MYA (Figure 2A) [36]. Previous research reported that the divergence time of *S. baicalensis* and *S. barbata* based on the *mat*K and *CHS* (chalcone synthase) genes was ∼3.35 MYA [37]. However, a genome-wide analysis identified 8 and 3 *CHS* genes in *S. baicalensis* and *S. barbata*, respectively, and the expansion and evolution of *CHS* negatively impacted the estimation of diversification history between these *Scutellaria* species.

Based on sequence homology, 17,265 orthologous gene pairs with synteny were identified between the *S. baicalensis* and *S. barbata* genomes, and the distribution of synonymous substitution rates (*Ks*) peaked at approximately 0.16, representing the speciation time of *S. baicalensis* and *S. barbata* (Figure 2B, Table S11). The mean *Ks* values of orthologous gene pairs with synteny and the divergence times among *S. baicalensis, S. barbata, S. miltiorrhiza, S. indicum*, and *Vitis vinifera* [38], showed the estimated synonymous substitutions per site per year as 1.30 ×10^−8^ for the test species (Table S11). In total, 7,812, 7,168, 6,984, and 7,711 paralogous gene pairs were identified, and the distribution of *Ks* values peaked at approximately 0.87, 0.86, 1.02 and 0.67 in *S. baicalensis, S. barbata, S. miltiorrhiza* and *S. indicum*, respectively (Figure 2B, Table S11). Based on the phylogenetic analysis, the WGD event happened before the divergence of *S. baicalensis, S. barbata, S. miltiorrhiza* and *S. indicum*. Then, we traced the divergence time of Lamiaceae and Pedaliaceae shared WGD event around 46.24-60.71 MYA (Table S11). The distribution of the *Ks* values of paralogous genes showed that no whole-genome duplication (WGD) events have occurred since the divergence of *S. miltiorrhiza, S. baicalensis* and *S. barbata*. Comparison of *S. baicalensis* and *S. barbata* genomes with an ancestral eudicot karyotype (AEK) genome [39], and with grape genome, also supported the structural rearrangement between *S. baicalensis* and *S. barbata* genomes, and the shared WGD event after WGT-γ event of grape (Figure 2C, Figure S9). The genome syntenic analysis indicated two copies of syntenic blocks from of Lamiaceae and Pedaliaceae species per corresponding grape block, which confirmed the recent WGD event before the divergence of *S. baicalensis, S. barbata, S. indicum* (Figure S10).

### Organ-specific localization of bioactive compounds

Baicalein, scutellarein, norwogonin, wogonin, and their glycosides (baicalin, scutellarin, norwogonoside and wogonoside) are the main bioactive compounds in *S. baicalensis* and *S. barbata*. We collected samples from the root, stem, leaf and flower tissues of *S. baicalensis* and *S. barbata* to detect the accumulation of active compounds. The results indicated that baicalein, norwogonin, wogonin, baicalin, norwogonoside and wogonoside mainly accumulated in the roots of *S. baicalensis* and *S. barbata*, while scutellarin was distributed in the aerial parts (stem, leaf and flower) of these species (Figure 1B, Figure S11, Table S12), providing a potential basis for the co-expression analysis of biosynthetic genes [23].

Transcriptome analysis of these four tissues from *S. baicalensis* and *S. barbata* included calculation of the FPKM values of 39,121 and 47,200 genes, respectively. Among them, 31.5% (12,320) and 40.6% (19,153) of the transcripts were not expressed (FPKM < 1) in any of the tested tissues. Based on k-means clustering, all the expressed genes from *S. baicalensis* and *S. barbata* were clustered into 48 groups (Figure S12 and S13). The expression levels of 3,421 genes from clusters 8, 20, 32, 33, 34, 39, and 47 in *S. baicalensis*, and 3,675 genes from clusters 2, 4, 21, 25, 27, 31, and 40 in *S. barbata* were significantly higher in the roots than in the other organs. The biosynthetic genes involved in the synthesis of *Scutellaria* specific flavones and glycosides, containing genes encoding chalcone synthase, chalcone isomerase, CYP450s, O-methyltransferase, glycosyltransferase and glycosyl hydrolases, were enriched, with high expression in the roots of *S. baicalensis* and *S. barbata* (Table S13 and S14).

### Conserved evolution of the chrysin and apigenin biosynthetic pathways in *S. baicalensis* and *S. barbata*

The main active compounds in the medicinal plants *S. baicalensis* and *S. barbata* are flavonoids, and the chrysin biosynthetic genes in *S. baicalensis* encoding 4-CoA ligase (4CL), chalcone synthase (CHS), chalcone isomerase (CHI), and flavone synthase (FNSII) have been cloned and functionally identified [14]. However, the gene locations, gene numbers and evolution of this pathway in the *S. baicalensis* and *S. barbata* genomes remain unclear. Here, we identified the same number of chrysin and apigenin biosynthetic genes in the *S. baicalensis* and *S. barbata* genomes and determined the expression levels of these genes, including phenylalanine ammonia lyase (PAL, 5 and 4), cinnamate 4-hydroxylase (C4H, 3 and 4), 4CL (9 and 14), CHS (8 and 3), CHI (1 and 1), and FNSII (3 and 3), in different tissues (**Figure 3**A, Table S15 and S16). Here, 18 orthologous gene pairs were found between the *S. baicalensis* and *S. barbata* genomes, and the *Ka*/*Ks* value (average 0.13) indicated purifying selection on flavone biosynthesis during evolution [40] (Figure 3B, Table S17). The PAL and CHS gene numbers in *S. baicalensis* were expanded compared to those in *S. barbata*; conversely, a significant duplication event of 4CL genes in *S. barbata* was found, suggesting that expansion via tandem duplication might have occurred after the separation of these *Scutellaria* species. The *Ks* values of 18 orthologous gene pairs of *S. baicalensis* and *S. barbata* in the chrysin and apigenin biosynthetic pathways indicated that the specific expansion of the SbaiPAL (SbaiPAL1 and SbaiPAL2), SbaiCHS (SbaiCHS2, SbaiCHS3, SbaiCHS4, and SbaiCHS5) and Sbar4CL (Sbar4CL1-1 and Sbar4CL1-2, Sbar4CL1-3 and Sbar4CL1-4, Sbar4CLL9-2 and Sbar4CLL9-3) genes had occurred via tandem duplication, after the speciation of *S. baicalensis* and *S. barbata* (Figure 3, Figure S14, Table S17).

**Figure 3.**
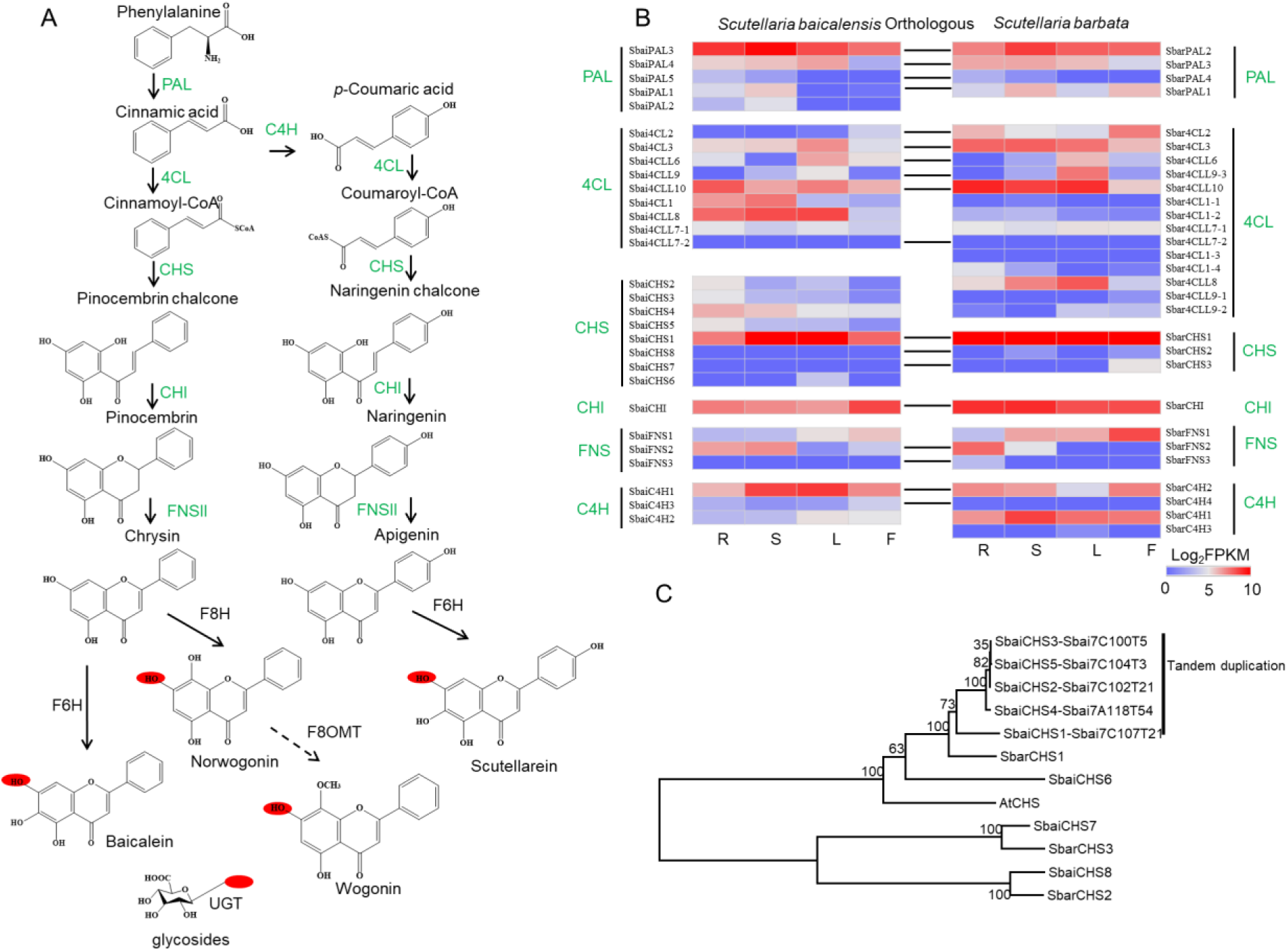
The potential biosynthetic pathway of flavonoids and species-specific gene expansion in *S. baicalensis* and *S. barbata*. A. Biosynthetic genes related to flavones and their glycosides. Phenylalanine ammonia lyase (PAL), cinnamate 4-hydroxylase (C4H), 4-CoA ligase (4CL), chalcone synthase (CHS), chalcone isomerase (CHI), flavone synthase (FNSII), flavone 6-hydroxylase (F6H) and flavone 8-hydroxylase (F8H). B. The expression profile and orthologous gene pairs of flavone biosynthetic genes in *S. baicalensis* and *S. barbata*. C. Tandem duplication and phylogenetic analysis of CHS genes.

Sbai4CLL7 and SbaiCHS1 have been reported to be related to the biosynthesis of specific 4’-deoxyflavones with cinnamic acid as a substrate in *S. baicalensis* [14]. Compared to *S. miltiorrhiza*, the 4CLL7 genes from the *Scutellaria* genus showed gene expansion, and the gene duplication of Sbai4CLL7-1 and Sbai4CLL7-2 occurred before the speciation of *S. baicalensis* and *S. barbata* (Figure S13). *Sbai4CLL7-1* and *Sbar4CLL7-1* showed no expression in the tested transcriptomes, and the duplication of the *Scutellaria*-specific 4CLL7-2 allowed the evolution of substrate preferences for the catalysis of cinnamic acid. The initial step and central hub for flavone biosynthesis is the catalysis of CHS; hence, the expression of CHS is required for the production of flavonoids, isoflavonoids, and other metabolites in plants [41]. Here, we also detected the highest expression levels of *SbaiCHS1* and *SbarCHS1* in all the tested samples; however, a recent expansion of CHS genes has occurred in *S. baicalensis*, and 4 additional paralogs of *SbaiCHS1* (*Sbai7C107T21*) were observed in chr7. Duplications of the *SbaiCHS2, SbaiCHS3, SbaiCHS4* and *SbaiCHS5* genes occurred after the speciation of *S. baicalensis* and *S. barbata* (Figure 3C). The nucleotide and amino acid sequences of SbaiCHS2 and SbaiCHS3 were identical, but SbaiCHS5 contained a variant K316 deletion. The divergence of *SbaiCHS1* and *SbarCHS1* occurred before the seperation of *S. miltiorrhiza* and the *Scutellaria* species, suggesting a conserved function of chalcone synthase in flavone biosynthesis. In addition, the tandemly duplicated SbaiCHS2-5 genes were more highly expressed in the roots of *S. baicalensis* than in other tissues (Figure 3C), suggesting that their species-specific evolution might be related to the biosynthesis of flavones and their glycosides, which are enriched in roots.

C4H is responsible for the biosynthesis of coumaroyl-CoA, which might be the restrictive precursor of the 4’-hydroxyl group involved in scutellarein biosynthesis. Here, we identified high expression of *SbaiC4H1* and *SbarC4H1* in the stems, leaves, and flowers of *S. baicalensis* and *S. barbata* (Figure 3B, Figure S14). This high expression level was positively correlated with the distribution of scutellarein, which is biosynthesized in the aerial parts of *S. baicalensis* and *S. barbata* (Figure 1B).

The SbaiFNSII2 gene, which has been reported to catalyze the formation of chrysin in *S. baicalensis*, presented high expression in the roots and stems, and its ortholog SbarFNSII2 was also significantly expressed in the roots of *S. barbata*. A genome collinearity analysis identified 566 orthologous gene pairs covering a region ∼6 Mb in length across chr3 of *S. baicalensis* and chr13 of *S. barbata*, including the tandem duplication of SbaiFNSII1-SbaiFNSII2 and SbarFNSII1-SbarFNSII2. This duplication occurred before the speciation of *S. baicalensis* and *S. barbata* (Figure S14). The majority of the FNSII region (∼85%) in *S. baicalensis* and *S. barbata* was assigned to the A compartment, indicating high transcriptional activity. The genome synteny of the FNSII region between *S. baicalensis* and *S. barbata* suggested the conserved evolution of flavone synthase.

### Functional divergence of flavone hydroxylase genes between *S. baicalensis* and *S. barbata*

CYP450 superfamily members, such as C4H (CYP73A family), FNSII (CYP93B family), flavone 6-hydroxylase (F6H, CYP82D family) and flavone 8-hydroxylase (F8H, CYP82D family), perform key modifications in flavone biosynthesis. SbaiCYP82D1 has been reported to have 6-hydroxylase activity on chrysin and apigenin to produce baicalein and scutellarein, respectively, and SbaiCYP82D2 can catalyze chrysin to norwogonin in *S. baicalensis* [15] (Figure S15). Here, we identified 418 and 398 CYP450 gene members, and 17 and 24 physical clusters of CYP450s (5 gene clusters per 500 kb) in the *S. baicalensis* and *S. barbata* genomes, respectively (Figure S16 and S17), suggesting a high frequency of CYP gene tandem duplication. Among them, 18 CYP82D members containing SbaiCYP82D1-9 and SbarCYP82D1-9 were identified in the *S. baicalensis* and *S. barbata* genomes; these genes might be responsible for the hydroxylation of chrysin and apigenin (Table S18). Consistent with a previous report, significant expression of *SbaiCYP82D1* and *SbaiCYP82D2* in the roots of *S. baicalensis* was detected, in accordance with the accumulation of baicalein, wogonin, and their glycosides (**Figure 4**A). However, *SbarCYP82D1* showed relatively high expression in stems and leaves, and *SbarCYP82D2* showed extremely low expression in all tissues of *S. barbata*, in contrast to the distributions of active flavones, suggesting a potential functional divergence of hydroxylation between *S. baicalensis* and *S. barbata*.

**Figure 4.**
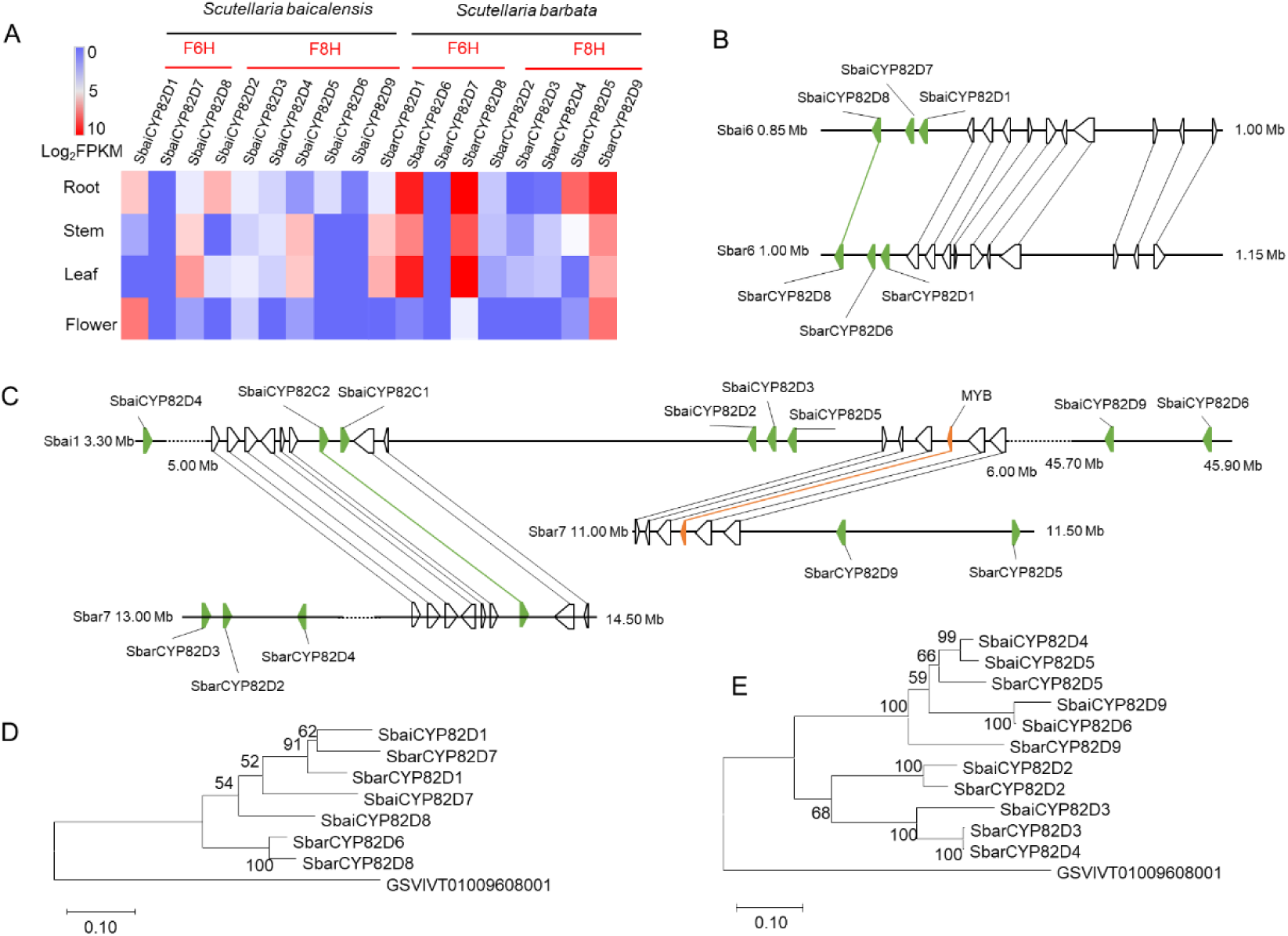
The tandem repeat of flavone hydroxylase genes in *S. baicalensis* and *S. barbata* revealed the divergent evolution. A. Identification and expression of CYP82D subfamily genes. Flavone 6-hydroxylase (F6H), and flavone 8-hydroxylase (F8H). B. Collinearity of CYP82D1 (F6H) regions between *S. baicalensis* and *S. barbata*. C. Synteny of CYP82D2 (F8H) regions between *S. baicalensis* and *S. barbata*. D. Phylogenetic tree of CYP82D1 groups. The grape CYP82D (GSVIVT01009608001) was chosen as outgroup. E. Phylogenetic tree of CYP82D2 groups. The grape CYP82D (GSVIVT01009608001) was chosen as outgroup.

Three-gene tandem duplications of *SbaiCYP82D1*-*SbaiCYP82D7*-*SbaiCYP82D8* and *SbarCYP82D1*-*SbarCYP82D6*-*SbarCYP82D8* (physical distance < 30 kb) on chr6 of *S. baicalensis* and *S. barbata* were identified (Figure 4B). According to the 150 kb collinearity analysis, 11 orthologous gene pairs, including CYP82D8 from *S. baicalensis* and *S. barbata*, presented conserved evolution. The phylogenetic analysis and *Ks* values of orthologous gene pairs indicated that the duplication of *SbarCYP82D8* and *SbarCYP82D6* occurred after the speciation of *S. barbata* (Table S19); however, duplication of *SbaiCYP82D8* and *SbaiCYP82D7* happened before the divergence of *S. baicalensis* and *S. barbata* (Figure 4D, Figure S18). This contradiction and evolutionary divergence supports the following proposed hypothesis: 1) the first duplication of *CYP82D8* produced the new *CYP82D1*, and the duplication event occurred around WGD event. 2) the second duplication of *CYP82D8* generated the new *CYP82D7*, similar to the tandem duplication of *SbaiCYP82D8*-*SbaiCYP82D7*-*SbaiCYP82D1* in *S. baicalensis*. 3) After speciation, the third duplication event of *SbarCYP82D8* uniquely occurred in the *S. barbata* genome and produced *SbarCYP82D6*; a recent gene transfer of *SbarCYP82D7* via transposon from chr6 to chr3 in *S. barbata* was predicted. An adjacent intact LTR/Gypsy in *SbarCYP82D7* was identified, and its insertion time was estimated to be ∼3.5 MYA. Given the evolution and high expression of *SbarCYP82D6* and *SbarCYP82D8*, we speculated that these two genes might be responsible for the F6H function in chrysin and apigenin synthesis *in vivo* in *S. barbata*.

The chromosome location of F8H functional members showed that *SbaiCYP82D2, SbaiCYP82D3, SbaiCYP82D4, SbaiCYP82D5, SbaiCYP82D6* and *SbaiCYP82D9* were distributed on chr1 of *S. baicalensis*, and *SbarCYP82D2, SbarCYP82D3, SbarCYP82D4, SbarCYP82D5* and *SbarCYP82D9* were located on chr7 of *S. barbata*. The structural rearrangement of large segments between chr1 of *S. baicalensis* and chr7 of *S. barbata* was found (Figure 4C, Figure S4). In addition, tandem duplications containing three CYP genes (*SbaiCYP82D2*-*SbaiCYP82D3*-*SbaiCYP82D5* and *SbarCYP82D3*-*SbarCYP82D2*-*SbarCYP82D4*) were identified (Figure 4C). The orthologous gene pairs (SbaiCYP82D2-SbarCYP82D2 and SbaiCYP82D3-SbarCYP82D3) presented high identity values of 90.11% and 83.72%. The duplications of *SbarCYP82D3*-*SbarCYP82D4, SbaiCYP82D4*-*SbaiCYP82D5*, and *SbaiCYP82D6*-*SbaiCYP82D9* occurred after the speciation of *S. baicalensis* and *S. barbata* (Table S19). However, the expression of *SbarCYP82D2, SbarCYP82D3* and *SbarCYP82D4* is slight in *S. barbata*, indicating functional divergence following species-specific duplication events. In contrast, the SbarCYP82D5 and SbarCYP82D9 were highly expressed in the roots of *S. barbata*, suggesting a potential F8H function in the biosynthesis of norwogonin.

## Conclusions

We reported two chromosome-level genomes of the medicinal plants *S. baicalensis* and *S. barbata* through the combination of second-generation sequencing (Illumina platform), third-generation sequencing (PacBio and Oxford Nanopore platforms), and Hi-C technologies. This study confirmed and traced the divergence times of *S. baicalensis* and *S. barbata*, which occurred 13.28 MYA, far earlier than previously reported. Comparative genomic analysis revealed similar TE proportions in the *S. baicalensis* and *S. barbata* genomes, while the recent LTR insertion in *S. barbata* might be an important factor resulting in chromosomal rearrangement and expansion. A WGD event (∼52.11-78.84 MYA) shared among *S. baicalensis, S. barbata, S. miltiorrhiza*, and *S. indicum*. The tandem duplication of paralogs after the speciation of *S. baicalensis* and *S. barbata* might be the most important contributor to the divergent evolution of flavonoid biosynthetic gene families, such as PAL, 4CL CHS, F6H and F8H. A determination of the distribution of flavone contents and transcriptome analysis supported the functional divergence of flavonoid biosynthetic genes between *S. baicalensis* and *S. barbata*. The two high-quality genomes reported in the present study will enrich genome research in the Lamiaceae and provide important insights for studies of breeding, evolution, chemodiversity and genome editing.

## Materials and methods

### Plant materials

*S. baicalensis* and *S. barbata* plants were cultivated in the experimental field of the IMPLAD (Institute of Medicinal Plant Development) (40°N and 116°E), Beijing, China. Four independent tissues from *S. baicalensis* and *S. barbata*, namely, root, stem, leaf, and flower tissues, were collected in three replicates. These tissues were used separately for the measurement of active compounds and RNA sequencing. High-quality DNA extracted from young leaves was used to construct libraries for Illumina, ONT and Sequel sequencing.

### Long-read sequencing and assemblies

The high-molecular-weight (HMW) genomic DNA of *S. baicalensis* and *S. barbata* was extracted in accordance with the method for megabase-sized DNA preparation [42]. HMW gDNA fragments (>20 kb) were selected using BluePippin. Long-read libraries were constructed following the protocols for the ONT (https://nanoporetech.com/) and PacBio Sequel platforms (https://www.pacb.com/). The ONT reads of *S. baicalensis* were generated using the ONT GridION X5 platform, and the library of *S. barbata* was sequenced using the Sequel platform. The raw ONT and SMRT reads were filtered via MinKNOW and SMRT Link, respectively. First, CANU (v1.7) was used to correct and trim the long reads from the ONT and Sequel platforms with the default parameters [43]. Then, the corrected and trimmed ONT and SMRT reads were assembled using SMARTdenovo (https://github.com/ruanjue/smartdenovo). Finally, Illumina short reads were used to polish the assembled contigs three times using Pilon (v1.22). The quality of the genome assemblies was estimated by a BUSCO (v2.0) search [44] and by mapping Illumina reads from the DNA and RNA libraries to the assembled genomes.

### Chromosome construction using Hi-C

Young leaves from *S. baicalensis* and *S. barbata* were fixed and crosslinked, and then, Hi-C libraries were constructed and sequenced using Illumina as described [33,34]. The short reads were mapped to the assembled genome using BWA [45], and the valid interaction pairs were selected using Hi-C Pro [46]. Then, the draft assemblies of *S. baicalensis* and *S. barbata* were anchored to chromosomes (2n = 18 and 2n = 26, respectively) using LACHESIS with the following parameters: CLUSTER MIN RE SITES = 62, CLUSTER MAX LINK DENSITY = 2, CLUSTER NONINFORMATIVE RATIO = 2, ORDER MIN N RES IN TRUN = 53, ORDER MIN N RES IN SHREDS = 52 [30].

### Genome annotation

The RepeatModeler (v1.0.9) package, including RECON and RepeatScout, was used to identify and classify the repeat elements of the *S. baicalensis* and *S. barbata* genomes. The repeat elements were then masked by RepeatMasker (v4.0.6). The long terminal repeat retrotransposons (LTR-RTs) in *S. baicalensis* and *S. barbata* were identified using LTR_Finder (v1.0.6) and LTR_retriever. Twenty-four samples from a total of eight different *S. baicalensis* and *S. barbata* tissues (roots, stems, leaves, and flowers) were subjected to RNA-Seq using the Illumina NovaSeq platform. The clean reads from *S. baicalensis* and *S. barbata* were *de novo* assembled using Trinity (v 2.2.0), and the coding regions in the assembled transcripts were predicted using TransDecoder (v2.1.0). The gene annotation of the masked *S. baicalensis* and *S. barbata* genome was *ab initio* predicted using the MAKER (v2.31.9) pipeline, integrating the assembled transcripts and protein sequences from *S. baicalensis, S. barbata*, and *A. thaliana* [47]. Noncoding RNAs and miRNAs were annotated by alignment to the Rfam and miRNA databases using INFERNAL (v1.1.2) and BLASTN, respectively. RNA-Seq reads from different *S. baicalensis* and *S. barbata* tissues were mapped to the masked genome using HISAT2 (v2.0.5), and the different expression levels of the annotated genes were calculated using Cufflinks (v2.2.1) [48].

### Genome evolution analysis

The full amino acid sequences of *S. baicalensis, S. barbata* and nin other angiosperms were aligned to orthologous groups using OrthoFinder [49]. The basal angiosperm *Amborella trichopoda*, was chosen as the outgroup. Single-copy genes were used to construct a phylogenetic tree using the RAxML package with PROTGAMMAJTT model and 1000 replicates (version 8.1.13). The divergence times among 11 plants were predicted using r8s program based on the estimated divergence times *Amborella_trichopoda*-*Vitis_vinifera* (173-199 MYA) and *Populus trichocarpa*-*Arabidopsis thaliana* (98-117 MYA). According to the phylogenetic analysis and divergence times, expansion and contraction of the gene families were identified using CAFÉ (v 3.1) [50]. The paralogous and orthologous gene pairs from *S. baicalensis, S. barbata*, and *S. miltiorrhiza* were identified, and the *Ka, Ks* and *Ka/Ks* values of *S. baicalensis*-*S. baicalensis, S. barbata*-*S. barbata, S. miltiorrhiza*-*S. miltiorrhiza, S. baicalensis*-*S. miltiorrhiza, S. baicalensis*-*S. barbata*, and *S. barbata*-*S. miltiorrhiza*, were calculated using the SynMap2 and DAGchainer method of CoGE Comparative Genomics Platform. The detection of synteny and collinearity among *S. baicalensis, S. barbata*, and *S. miltiorrhiza* was performed using MCscan X(v1.1) [51].

### Identification of gene families related to flavone biosynthesis

Protein sequences of the PAL, 4CL, C4H, CHS, CHI, and FNSII gene family members in *A. thaliana* were downloaded from the TAIR database, and F6H and F8H in *S. baicalensis* were obtained from a previous study. Then, these sequences were searched against the *S. baicalensis* and *S. barbata* protein sequences using BLASTP with an E value cutoff of 1e-10. The conserved domains of the protein sequences of candidate genes were further searched in the Pfam database using hidden Markov models [52]. Full-length protein sequences were used to construct phylogenetic trees using the maximum likelihood method with the Jones-Taylor-Thornton (JTT) model and 1,000 bootstrap replicates [53]. A detailed description of some materials and methods used is provided in Supplementary methods and results.

## Data availability

The raw sequence data reported in this paper have been deposited in the Genome Sequence Archive [54] in BIG Data Center [55], Beijing Institute of Genomics (BIG), Chinese Academy of Sciences, under accession numbers CRA001730 that are publicly accessible at http://bigd.big.ac.cn/gsa. The assembled genomes and gene structures were also submitted to CoGe with id54175 for *S. baicalensis* and id54176 for *S. barbata*.

## Authors’ Contributions

ZX and JS designed and coordinated the study. ZX assembled and analyzed the genome. RX, JW, SZ, YZ, and JC supplied plant materials. RG, XP, and CH performed the experiments and analyzed the data. ZX, RG, and JS wrote and edited the manuscript.

## Acknowledgments

This work was supported by the National Natural Science Foundation of China (Grant No. 31700264) and the Chinese Academy of Medical Sciences (CAMS) Innovation Fund for Medical Sciences (CIFMS) (Grant No. 2016-I2M-3-016).

## Supplementary material

Supplementary Figure S1. Genome size estimation using flow cytometry and the 21 *k*-mer distribution. A. Flow cytometry analysis using *Salvia miltiorrhiza* data as internal standards. B. The 21 *k*-mer distribution from Illumina short reads of *S. baicalensis* and *S. barbata*.

Supplementary Figure S2. Hi-C intrachromosomal contact map of *S. baicalensis* and *S. barbata* chromosomes. The red diagonal line indicates a high number of intrachromosomal contacts. A. Hi-C heatmap of *S. baicalensis*. B. Hi-C heatmap of *S. barbata*.

Supplementary Figure S3. Genome synteny analysis of *S. baicalensis* and *S. barbata* using MCscanX.

Supplementary Figure S4. The alignment of large-scale DNA sequences between *S. baicalensis* and *S. barbata* using MUMmer.

Supplementary Figure S5. Insertion time distribution of intact LTR-RTs in *S. baicalensis* and *S. barbata* assuming a mutation rate of *μ*=1.3×10^−8^ (per bp per year).

Supplementary Figure S6. Genome-wide chromatin packing analysis in *S. baicalensis*. A. The intrachromosomal interactions revealing the A/B compartments of *S. baicalensis*. B. The ratio of TE and gene numbers between the A and B compartments. C. The interchromosomal interactions of *S. baicalensis*.

Supplementary Figure S7. Genome-wide chromatin packing analysis in *S. barbata*. A. The intrachromosomal interactions revealing the A/B compartments of *S. barbata*. B. The ratio of TE and gene numbers between the A and B compartments. C. The interchromosomal interactions of *S. barbata*.

Supplementary Figure S8. The Gene family expansion and contraction of candidate species. The number of expansion and contraction events of 20 nodes are listed in Table S10.

Supplementary Figure S9. The grape genome was painted into *S. baicalensis* and *S. barbata* genome, respectively. The synteny from paralogs was detected by MCScanX.

Supplementary Figure S10. The gene syntenic analysis within candidate species. Dot plot presented that the gene synteny of grape-Sesame, grape-*S. baicalensis*, and grape-*S. barbata*, respectively. The red circles highlighted the duplication events after WGD-γ event.

Supplementary Figure S11. Ultraperformance liduid chromatography (UPLC) detection (280 nm) of flavonoid contents. The UPLC detection of flavonoids in different tissues of *S. baicalensis* and *S. barbata*, including baicalein, scutellarein, wogonin, and their glycosides (baicalin, scutellarin, and wogonoside). The compound information, including detailed retention times and spectrum data, is listed in Table S12. A. Flavonoid contents of *S. baicalensis*. B. Flavonoid contents of *S. barbata*.

Supplementary Figure S12. Gene expression clusters based on *k*-means in *S. baicalensis*. All expressed genes were clustered into 48 clusters in different *S. baicalensis* tissues, namely, root, stem, leaf, and flower tissues.

Supplementary Figure S13. Gene expression clusters based on *k*-means in *S. barbata*. All expressed genes were clustered into 48 clusters in different *S. barbata* tissues, namely, root, stem, leaf, and flower tissues.

Supplementary Figure S14. Phylogenetic analysis of PAL, C4H, 4CL, and FNSII from *S. baicalensis* and *S. barbata* using the maximum likelihood method.

Supplementary Figure S15. The potential biosynthetic pathway of baicalein, scutellarein, wogonin, and their glycosides (baicalin, scutellarin, and wogonoside), catalyzing chrysin and apigenin.

Supplementary Figure S16. The physical clusters of CYP450s (5 gene clusters per 500 kb) in *S. baicalensis*.

Supplementary Figure S17. The physical clusters of CYP450s (5 gene clusters per 500 kb) in *S. barbata*.

Supplementary Figure S18. The phylogenetic analysis of CYP82D, CYP93B, and CYP73A members from *S. baicalensis* and *S. barbata* using the maximum likelihood method.

Supplementary Table S1. The statistics of sequencing data from the SMRT and ONT platforms and corrected reads using CANU.

Supplementary Table S2. The assembled statistics of the *S. baicalensis* and *S. barbata* genome.

Supplementary Table S3. The genome synteny between *S. baicalensis* and *S. barbata*.

Supplementary Table S4. Genome annotations among *S. baicalensis, S. barbata* and *S. miltiorrhiza*.

Supplementary Table S5. Annotation of *S. baicalensis* TEs.

Supplementary Table S6. Annotation of *S. barbata* TEs.

Supplementary Table S7. Summary of intact LTR retrotransposons in *S. baicalensis* and *S. barbata*.

Supplementary Table S8. Annotation of *S. baicalensis* and *S. barbata* rRNA.

Supplementary Table S9. Identification of SSRs in the *S. baicalensis* and *S. barbata* genome.

Supplementary Table S10. Gene family expansion and contraction of candidate species according to phylogenetic analysis (*P* < 0.01).

Supplementary Table S11. The *Ks* value and divergence time of paralogous or orthologous gene pairs among *S. baicalensis, S. barbata, S. miltiorrhiza, S. indicum*, and *V. vinifera*.

Supplementary Table S12. The compound information of UPLC detection including retention time and spectrum, is shown in Supplementary Figure S11.

Supplementary Table S13. The Pfam annotation of genes with high expression in the roots of *S. baicalensis*.

Supplementary Table S14. The Pfam annotation of genes with high expression in the roots of *S. barbata*.

Supplementary Table S15. The expression of chrysin and apigenin biosynthetic genes in different organs of *S. baicalensis*.

Supplementary Table S16. The expression of chrysin and apigenin biosynthetic genes in different organs of *S. barbata*.

Supplementary Table S17. The *Ka* and *Ks* analysis of chrysin and apigenin biosynthetic genes in *S. baicalensis* and *S. barbata*.

Supplementary Table S18. The expression of CYP82D members in different tissues of *S. baicalensis* and *S. barbata*.

Supplementary Table S19. The *Ks* values of gene pairs related to flavone biosynthesis in *S. baicalensis* and *S. barbata*.

## References

[1] Winkel-Shirley B. Flavonoid biosynthesis. A colorful model for genetics, biochemistry, cell biology, and biotechnology. Plant Physiol 2001; 126: 485–93.

[2] Winkel-Shirley B. Biosynthesis of flavonoids and effects of stress. Curr Opin Plant Biol 2002; 5: 218–23.

[3] Grotewold E. The genetics and biochemistry of floral pigments. Annu Rev Plant Biol 2006; 57: 761–80.

[4] Shang X, He X, He X, Li M, Zhang R, Fan P, et al. The genus *Scutellaria* an ethnopharmacological and phytochemical review. J Ethnopharmacol 2010; 128: 279–313.

[5] Grzegorczyk-Karolak I, Wiktorek-Smagur A, Hnatuszko-Konka K. An untapped resource in the spotlight of medicinal biotechnology: the genus *Scutellaria*. Curr Pharm Biotechnol 2018; 19: 358–71.

[6] Chinese Pharmacopoeia Commission. Pharmacopoeia of the People’s Republic of China. Beijing: China Medical Science Press; 2015.

[7] Zhang Z, Lian XY, Li S, Stringer JL. Characterization of chemical ingredients and anticonvulsant activity of American skullcap (*Scutellaria lateriflora*). Phytomedicine 2009; 16: 485–93.

[8] Qiao X, Li R, Song W, Miao WJ, Liu J, Chen HB, et al. A targeted strategy to analyze untargeted mass spectral data: Rapid chemical profiling of *Scutellaria baicalensis* using ultra-high performance liquid chromatography coupled with hybrid quadrupole orbitrap mass spectrometry and key ion filtering. J Chromatogr A 2016; 1441: 83–95.

[9] Yan B, Xu W, Su S, Zhu S, Zhu Z, Zeng H, et al. Comparative analysis of 15 chemical constituents in *Scutellaria baicalensis* stem-leaf from different regions in China by ultra-high performance liquid chromatography with triple quadrupole tandem mass spectrometry. J Sep Sci 2017; 40: 3570–81.

[10] Zhao Q, Chen XY, Martin C. *Scutellaria baicalensis*, the golden herb from the garden of Chinese medicinal plants. Sci Bull (Beijing) 2016; 61: 1391–8.

[11] Dai J, Liang K, Zhao S, Jia W, Liu Y, Wu H, et al. Chemoproteomics reveals baicalin activates hepatic CPT1 to ameliorate diet-induced obesity and hepatic steatosis. Proc Natl Acad Sci U S A 2018; 115: E5896–905.

[12] Guo HX, Liu DH, Ma Y, Liu JF, Wang Y, Du ZY, et al. Long-term baicalin administration ameliorates metabolic disorders and hepatic steatosis in rats given a high-fat diet. Acta Pharmacol Sin 2009; 30: 1505–12.

[13] Chen S, Song J, Sun C, Xu J, Zhu Y, Verpoorte R, et al. Herbal genomics: examining the biology of traditional medicines. Science 2015; 347: S27–8.

[14] Zhao Q, Zhang Y, Wang G, Hill L, Weng JK, Chen XY, et al. A specialized flavone biosynthetic pathway has evolved in the medicinal plant, *Scutellaria baicalensis*. Sci Adv 2016; 2: e1501780.

[15] Zhao Q, Cui MY, Levsh O, Yang D, Liu J, Li J, et al. Two CYP82D enzymes function as flavone hydroxylases in the biosynthesis of root-specific 4’-deoxyflavones in *Scutellaria baicalensis*. Mol Plant 2018; 11: 135–48.

[16] Liu X, Cheng J, Zhang G, Ding W, Duan L, Yang J, et al. Engineering yeast for the production of breviscapine by genomic analysis and synthetic biology approaches. Nat Commun 2018; 9: 448.

[17] Li JH, Tian CF, Xia YH, Mutanda I, Wang KB, Wang Y. Production of plant-specific flavones baicalein and scutellarein in an engineered *E. coli* from available phenylalanine and tyrosine. Metab Eng 2019; 52: 124–33.

[18] Xu Z, Song J. The 2-oxoglutarate-dependent dioxygenase superfamily participates in tanshinone production in *Salvia miltiorrhiza*. J Exp Bot 2017; 68: 2299–308.

[19] Cao W, Wang Y, Shi M, Hao X, Zhao W, Wang Y, et al. Transcription factor SmWRKY1 positively promotes the biosynthesis of tanshinones in *Salvia miltiorrhiza*. Front Plant Sci 2018; 9: 554.

[20] Huang Q, Sun M, Yuan T, Wang Y, Shi M, Lu S, et al. The AP2/ERF transcription factor SmERF1L1 regulates the biosynthesis of tanshinones and phenolic acids in *Salvia miltiorrhiza*. Food Chem 2019; 274: 368–75.

[21] Sun M, Shi M, Wang Y, Huang Q, Yuan T, Wang Q, et al. The biosynthesis of phenolic acids is positively regulated by the JA-responsive transcription factor ERF115 in *Salvia miltiorrhiza*. J Exp Bot 2019; 70: 243–54.

[22] Xu H, Song J, Luo H, Zhang Y, Li Q, Zhu Y, et al. Analysis of the genome sequence of the medicinal plant *Salvia miltiorrhiza*. Mol Plant 2016; 9: 949–52.

[23] Xu Z, Peters RJ, Weirather J, Luo H, Liao B, Zhang X, et al. Full-length transcriptome sequences and splice variants obtained by a combination of sequencing platforms applied to different root tissues of *Salvia miltiorrhiza* and tanshinone biosynthesis. Plant J 2015; 82: 951–61.

[24] Xin T, Zhang Y, Pu X, Gao R, Xu Z, Song J. Trends in Herbgenomics. Sci China Life Sci 2019; 62: 288–308

[25] Xu Z, Xin T, Bartels D, Li Y, Gu W, Yao H, et al. Genome analysis of the ancient tracheophyte *Selaginella tamariscina* reveals evolutionary features relevant to the acquisition of desiccation tolerance. Mol Plant 2018; 11: 983–94

[26] Schmidt MH, Vogel A, Denton AK, Istace B, Wormit A, van de Geest H, et al. *De novo* assembly of a new *Solanum pennellii* accession using nanopore sequencing. Plant Cell 2017; 29: 2336–48.

[27] Guo L, Winzer T, Yang X, Li Y, Ning Z, He Z, et al. The opium poppy genome and morphinan production. Science 2018; 362: 343–7.

[28] Xu H, Song J, Luo H, Zhang Y, Li Q, Zhu Y, et al. Analysis of the genome sequence of the medicinal plant *Salvia miltiorrhiza*. Mol Plant 2016; 9: 949–52.

[29] Vining KJ, Johnson SR, Ahkami A, Lange I, Parrish AN, Trapp SC, et al. Draft genome sequence of mentha longifolia and development of resources for mint cultivar improvement. Mol Plant 2017; 10: 323–39.

[30] Burton JN, Adey A, Patwardhan RP, Qiu R, Kitzman JO, Shendure J. Chromosome-scale scaffolding of *de novo* genome assemblies based on chromatin interactions. Nat Biotechnol 2013; 31: 1119–25.

[31] VanBuren R, Wai CM, Ou S, Pardo J, Bryant D, Jiang N, et al. Extreme haplotype variation in the desiccation-tolerant clubmoss *Selaginella lepidophylla*. Nat Commun 2018; 9: 13.

[32] Bennetzen JL, Wang H. The contributions of transposable elements to the structure, function, and evolution of plant genomes. Annu Rev Plant Biol 2014; 65: 505–30.

[33] Liu C, Cheng YJ, Wang JW, Weigel D. Prominent topologically associated domains differentiate global chromatin packing in rice from *Arabidopsis*. Nat Plants 2017; 3: 742–8.

[34] Wang C, Liu C, Roqueiro D, Grimm D, Schwab R, Becker C, et al. Genome-wide analysis of local chromatin packing in *Arabidopsis thaliana*. Genome Res 2015; 25: 246–56.

[35] Song C, Liu Y, Song A, Dong G, Zhao H, Sun W, et al. The *Chrysanthemum nankingense* genome provides insights into the evolution and diversification of chrysanthemum flowers and medicinal traits. Mol Plant 2018; 11: 1482–91.

[36] Wang L, Yu S, Tong C, Zhao Y, Liu Y, Song C, et al. Genome sequencing of the high oil crop sesame provides insight into oil biosynthesis. Genome Biol 2014; 15: R39.

[37] Chiang YC, Huang BH, Liao PC. Diversification, biogeographic pattern, and demographic history of Taiwanese *Scutellaria* species inferred from nuclear and chloroplast DNA. PLoS One 2012; 7: e50844.

[38] Jaillon O, Aury JM, Noel B, Policriti A, Clepet C, Casagrande A, et al. The grapevine genome sequence suggests ancestral hexaploidization in major angiosperm phyla. Nature 2007; 449: 463–7.

[39] Murat F, Armero A, Pont C, Klopp C, Salse J. Reconstructing the genome of the most recent common ancestor of flowering plants. Nat Genet 2017; 49: 490–6.

[40] Navarro A, Barton NH. Chromosomal speciation and molecular divergence– accelerated evolution in rearranged chromosomes. Science 2003; 300: 321–4.

[41] Zhang X, Abrahan C, Colquhoun TA, Liu CJ. A proteolytic regulator controlling chalcone synthase stability and flavonoid biosynthesis in *Arabidopsis*. Plant Cell 2017; 29: 1157–74.

[42] Zhang M, Zhang Y, Scheuring CF, Wu CC, Dong JJ, Zhang HB. Preparation of megabase-sized DNA from a variety of organisms using the nuclei method for advanced genomics research. Nat Protoc 2012; 7: 467–78.

[43] Koren S, Walenz BP, Berlin K, Miller JR, Bergman NH, Phillippy AM. Canu: scalable and accurate long-read assembly via adaptive *k*-mer weighting and repeat separation. Genome Res 2017; 27: 722–36.

[44] Simao FA, Waterhouse RM, Ioannidis P, Kriventseva EV, Zdobnov EM. BUSCO: assessing genome assembly and annotation completeness with single-copy orthologs. Bioinformatics 2015; 31: 3210–2.

[45] Li H, Durbin R. Fast and accurate short read alignment with Burrows-Wheeler transform. Bioinformatics 2009; 25: 1754–60.

[46] Servant N, Varoquaux N, Lajoie BR, Viara E, Chen CJ, Vert JP, et al. HiC-Pro: an optimized and flexible pipeline for Hi-C data processing. Genome Biol 2015; 16: 259.

[47] Cantarel BL, Korf I, Robb SM, Parra G, Ross E, Moore B, et al. MAKER: an easy-to-use annotation pipeline designed for emerging model organism genomes. Genome Res 2008; 18: 188–96.

[48] Ghosh S, Chan CK. Analysis of RNA-seq data using TopHat and Cufflinks. Methods Mol Biol 2016; 1374: 339–61.

[49] Emms DM, Kelly S. OrthoFinder: phylogenetic orthology inference for comparative genomics. Gonome Biol 2019; 20: 238.

[50] De Bie T, Cristianini N, Demuth JP, Hahn MW. CAFE: a computational tool for the study of gene family evolution. Bioinformatics 2006; 22: 1269–71.

[51] Wang Y, Tang H, Debarry JD, Tan X, Li J, Wang X, et al. MCScanX: a toolkit for detection and evolutionary analysis of gene synteny and collinearity. Nucleic Acids Res 2012; 40: e49.

[52] El-Gebali S, Mistry J, Bateman A, Eddy SR, Luciani A, Potter SC, et al. The Pfam protein families database in 2019. Nucleic Acids Res 2019; 47: D427–32.

[53] Kumar S, Stecher G, Li M, Knyaz C, Tamura K. MEGA X: molecular evolutionary genetics analysis across computing platforms. Mol Biol Evol 2018; 35: 1547–9.

[54] Wang Y, Song F, Zhu J, Zhang S, Yang Y, Chen T, et al. GSA: genome sequence archive. Genomics Proteomics Bioinformatics 2017; 15: 14–8.

[55] Zhang Z, Zhao W, Xiao J, Bao Y, Wang F, Hao L, et al. Database Resources of the BIG Data Center in 2019. Nucleic Acids Res 2019; 47: D8–14.

