## Supplemental Figure 1 for "Comparative Genome Analysis of *Scutellaria baicalensis* and *Scutellaria barbata* Reveals the Evolution of Active Flavonoid Biosynthesis"

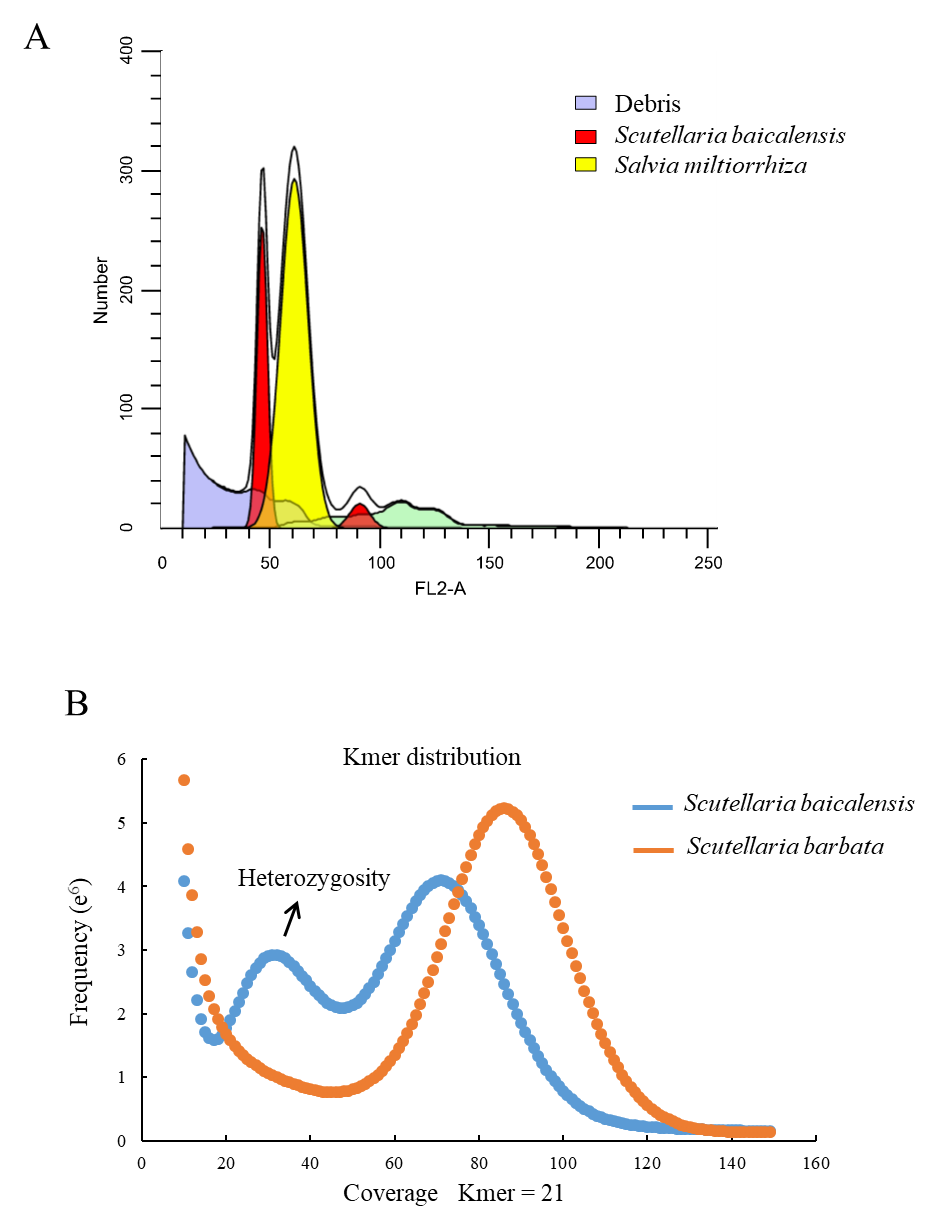

Supplementary Figure S1. Genome size estimation using flow cytometry and the 21 *k*-mer distribution. A. Flow cytometry analysis using *Salvia miltiorrhiza* data as internal standards. B. The 21 *k*-mer distribution from Illumina short reads of *S. baicalensis* and *S. barbata*.
