## Supplemental Figure 2 for "Comparative Genome Analysis of *Scutellaria baicalensis* and *Scutellaria barbata* Reveals the Evolution of Active Flavonoid Biosynthesis"

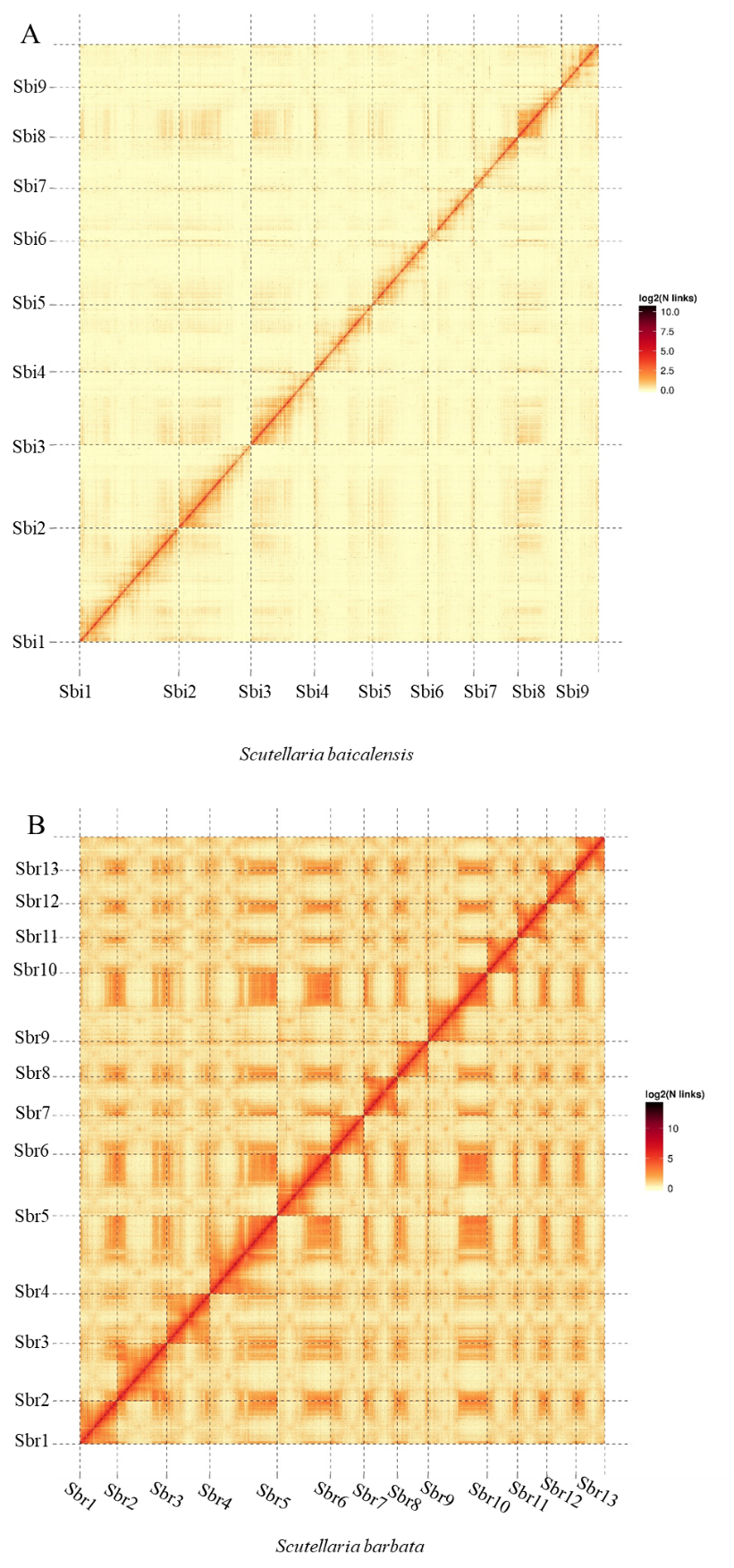

Supplementary Figure S2. Hi-C intrachromosomal contact map of *S. baicalensis* and *S. barbata* chromosomes. The red diagonal line indicates a high number of intrachromosomal contacts. A. Hi-C heatmap of *S. baicalensis*. B. Hi-C heatmap of *S. barbata*.
