## Supplemental Figure 3 for "Comparative Genome Analysis of *Scutellaria baicalensis* and *Scutellaria barbata* Reveals the Evolution of Active Flavonoid Biosynthesis"

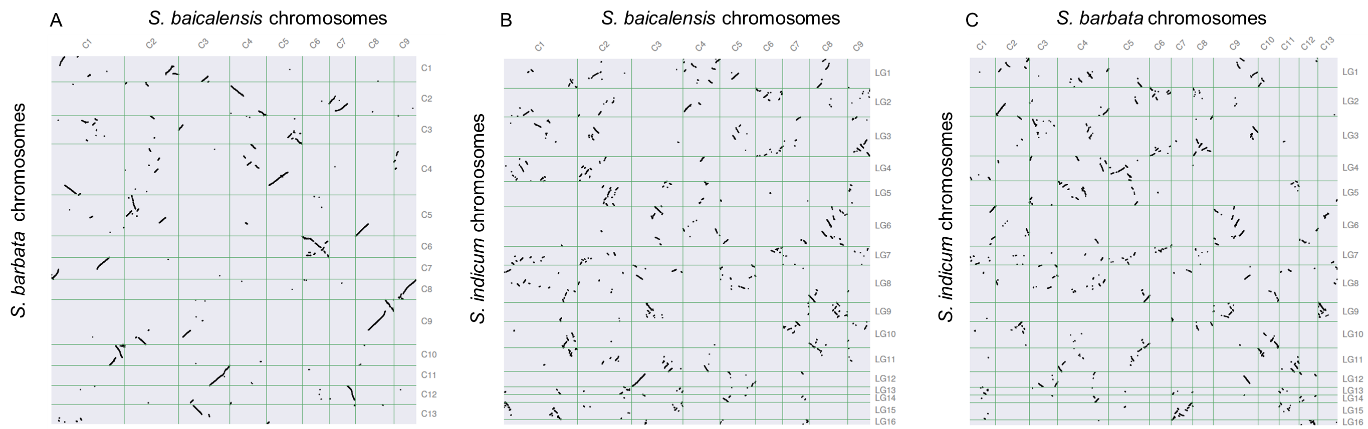

Supplementary Fig. S3. Genome synteny analysis between *S. baicalensis* and *S. barbata* (A), *S. baicalensis* and sesame (B), *S. barbata* and sesame (C), respectively, using MCScanX.
