## Supplemental Figure 4 for "Comparative Genome Analysis of *Scutellaria baicalensis* and *Scutellaria barbata* Reveals the Evolution of Active Flavonoid Biosynthesis"

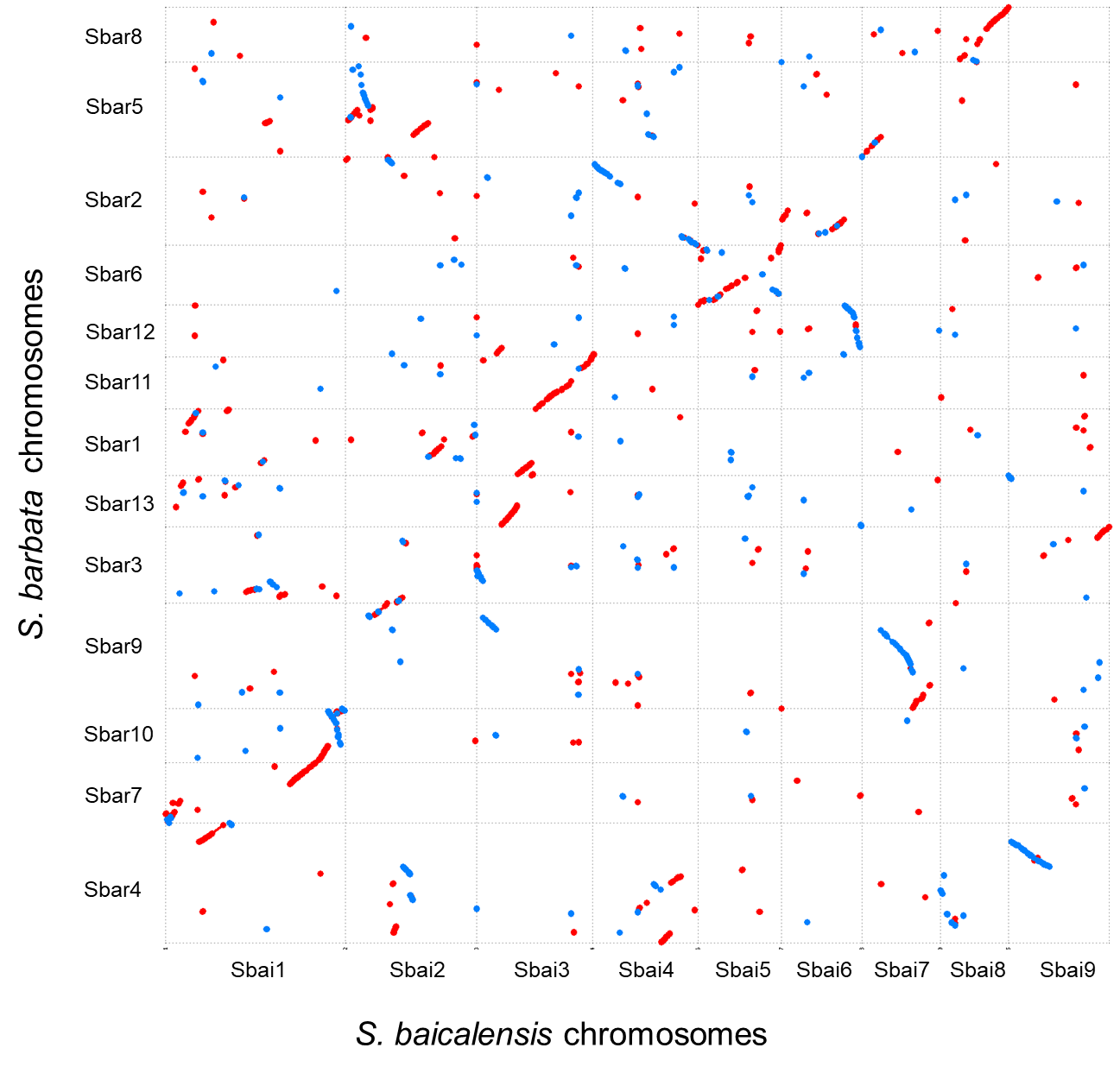

Supplementary Figure S4. The alignment of large-scale DNA sequences between *S. baicalensis* and *S. barbata* with the minimum mapping length of 100 kb.
