## Supplemental Figure 5 for "Comparative Genome Analysis of *Scutellaria baicalensis* and *Scutellaria barbata* Reveals the Evolution of Active Flavonoid Biosynthesis"

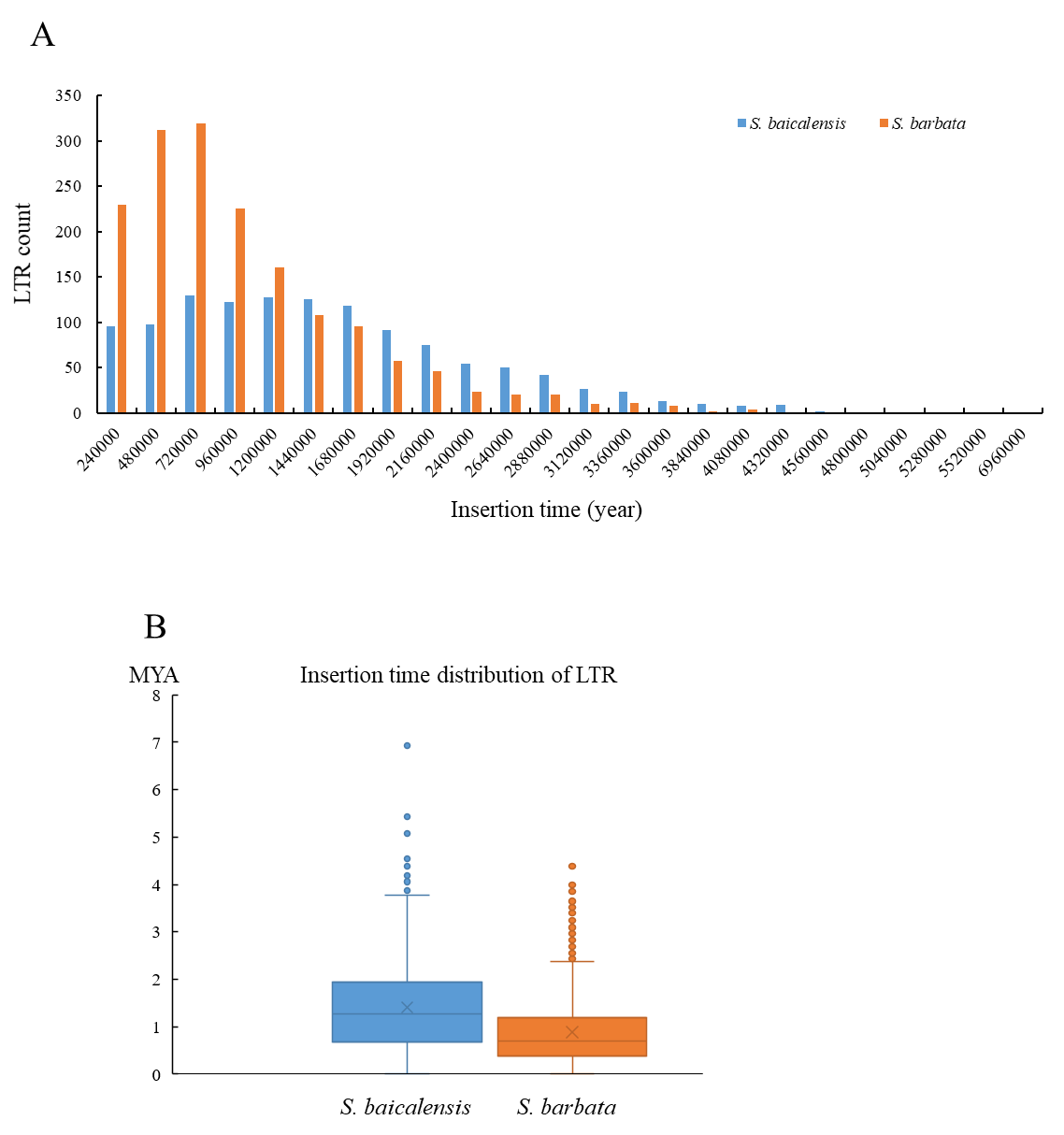

Supplementary Figure S5. Insertion time distribution of intact LTR-RTs in *S. baicalensis* and *S. barbata* assuming a mutation rate of *μ*=1.3×10^-8^ (per bp per year).
