## Supplemental Figure 6 for "Comparative Genome Analysis of *Scutellaria baicalensis* and *Scutellaria barbata* Reveals the Evolution of Active Flavonoid Biosynthesis"

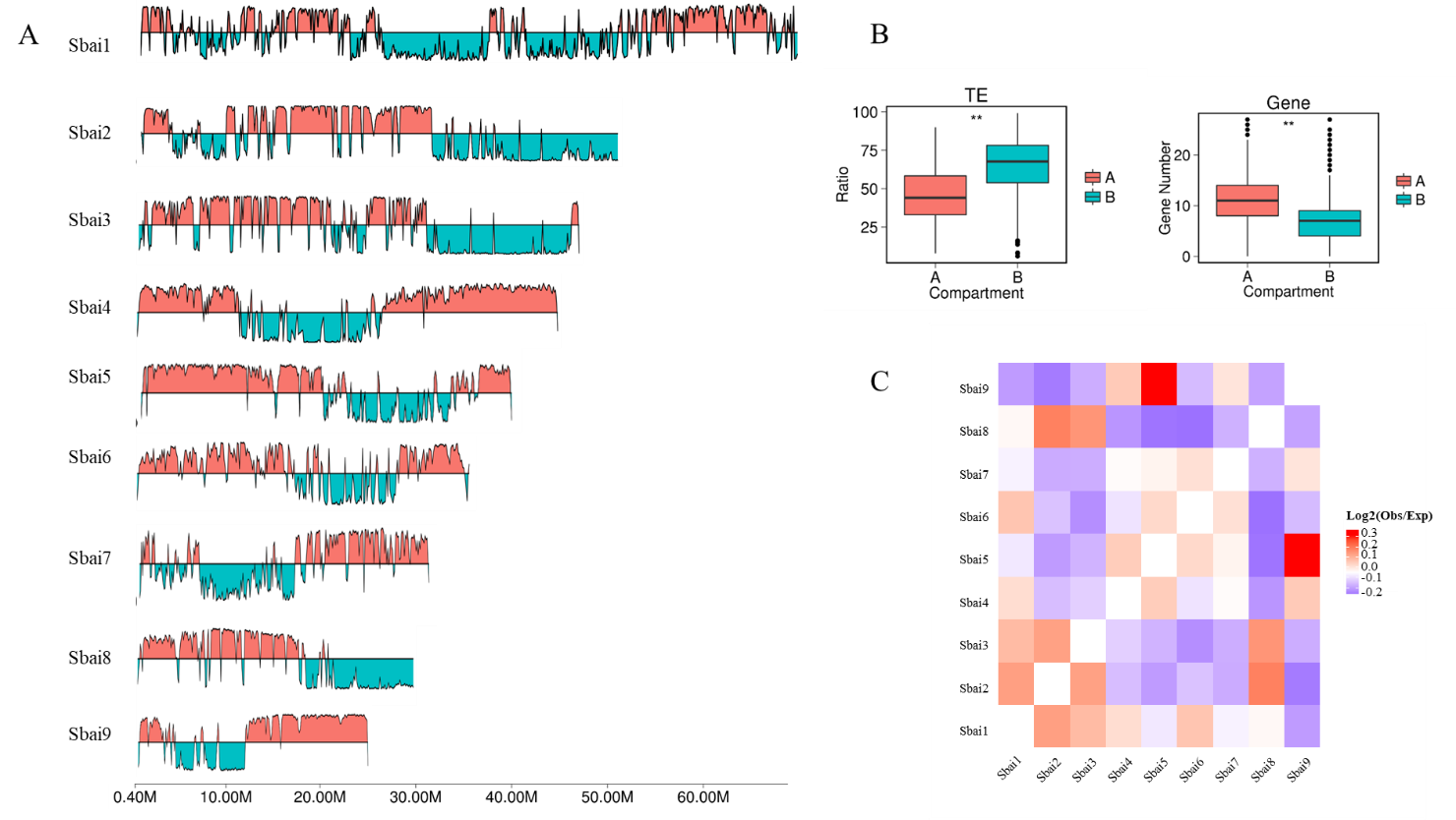

Supplementary Figure S6. Genome-wide chromatin packing analysis in *S. baicalensis*. A. The intrachromosomal interactions revealing the A/B compartments of *S. baicalensis*. B. The ratio of TE and gene numbers between the A and B compartments. C. The interchromosomal interactions of *S. baicalensis*.
