## Supplemental Figure 8 for "Comparative Genome Analysis of *Scutellaria baicalensis* and *Scutellaria barbata* Reveals the Evolution of Active Flavonoid Biosynthesis"

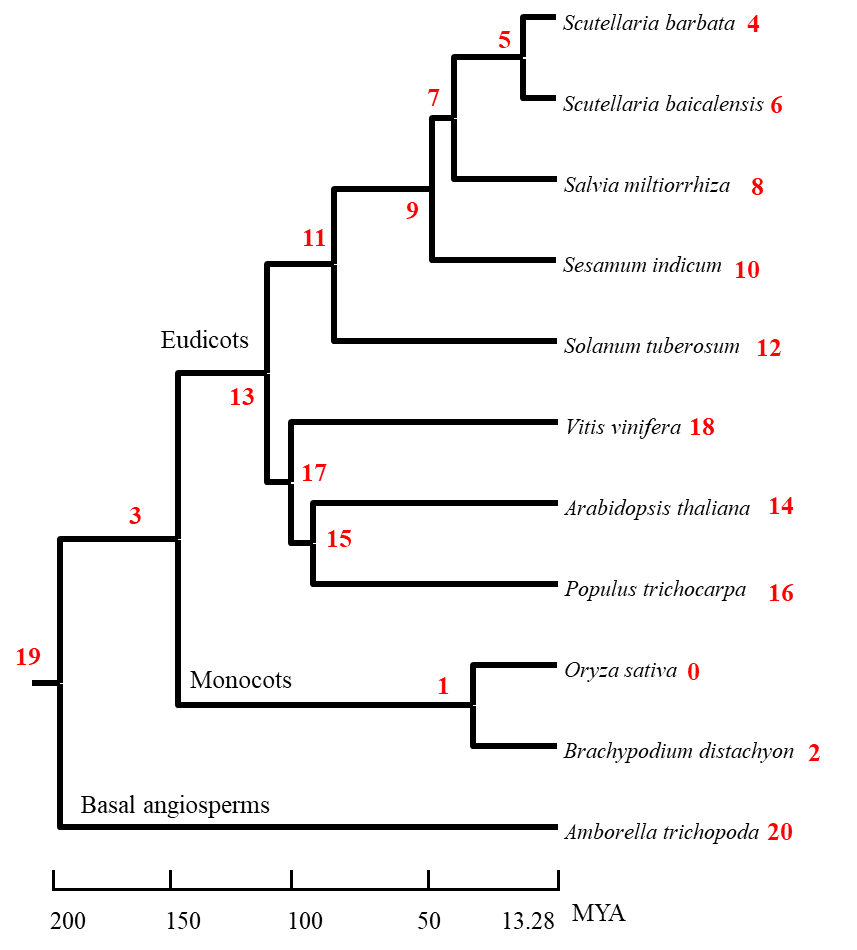

Supplementary Figure S8. The Gene family expansion and contraction of candidate species according to phylogenetic analysis (*P* < 0.01). The number of expansion and contraction events of 20 nodes are listed in Table S10.
