## Supplemental Figure 9 for "Comparative Genome Analysis of *Scutellaria baicalensis* and *Scutellaria barbata* Reveals the Evolution of Active Flavonoid Biosynthesis"

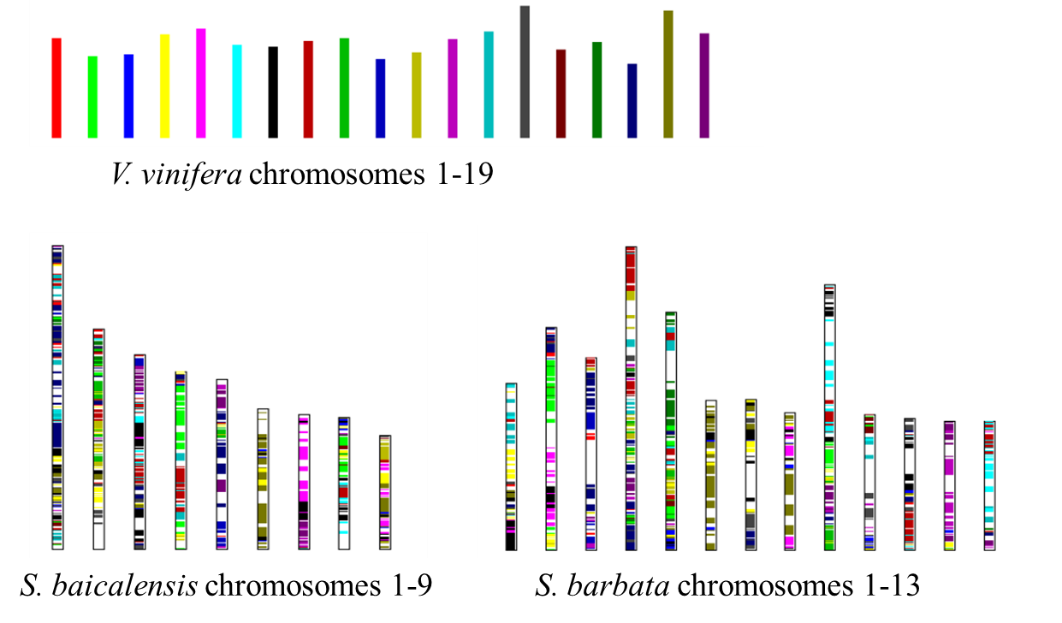

Supplementary Figure S9. The grape genome were painted into *S. baicalensis* and *S. barbata* genome, respectively, based on the gene collinearity using MCScanX.
