## Supplemental Figure 10 for "Comparative Genome Analysis of *Scutellaria baicalensis* and *Scutellaria barbata* Reveals the Evolution of Active Flavonoid Biosynthesis"

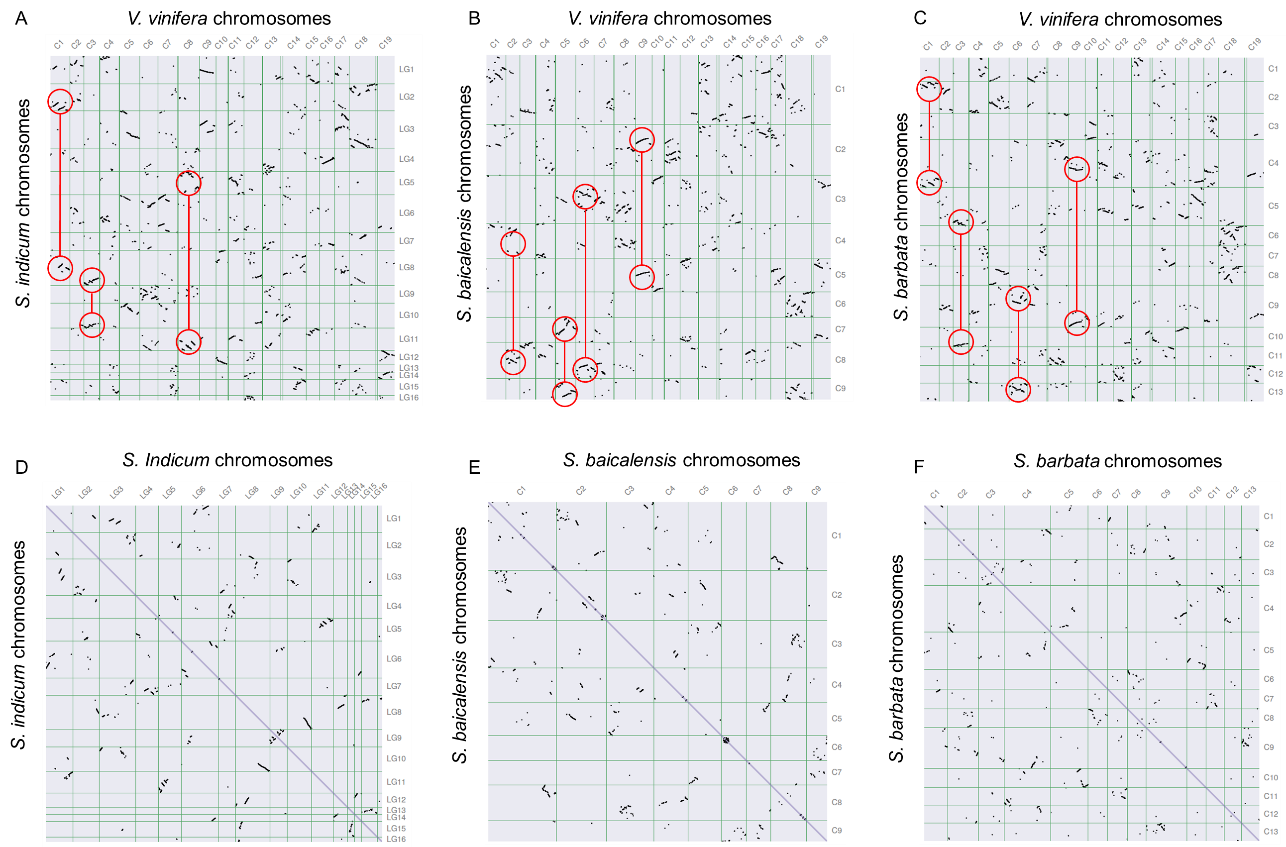

Supplementary Figure S10. Dot plots presented that the gene synteny between grape and Sesame (A), grape and *S. baicalensis*(B), grape and *S. barbata* (C), respectively. The red circles highlighted the duplication events after WGD-γ event. Dot plots of paralogs in Sesame (D), *S. baicalensis* (E), and *S. barbata* (F) to show the potential duplication events.
