## Supplemental Figure 11 for "Comparative Genome Analysis of *Scutellaria baicalensis* and *Scutellaria barbata* Reveals the Evolution of Active Flavonoid Biosynthesis"

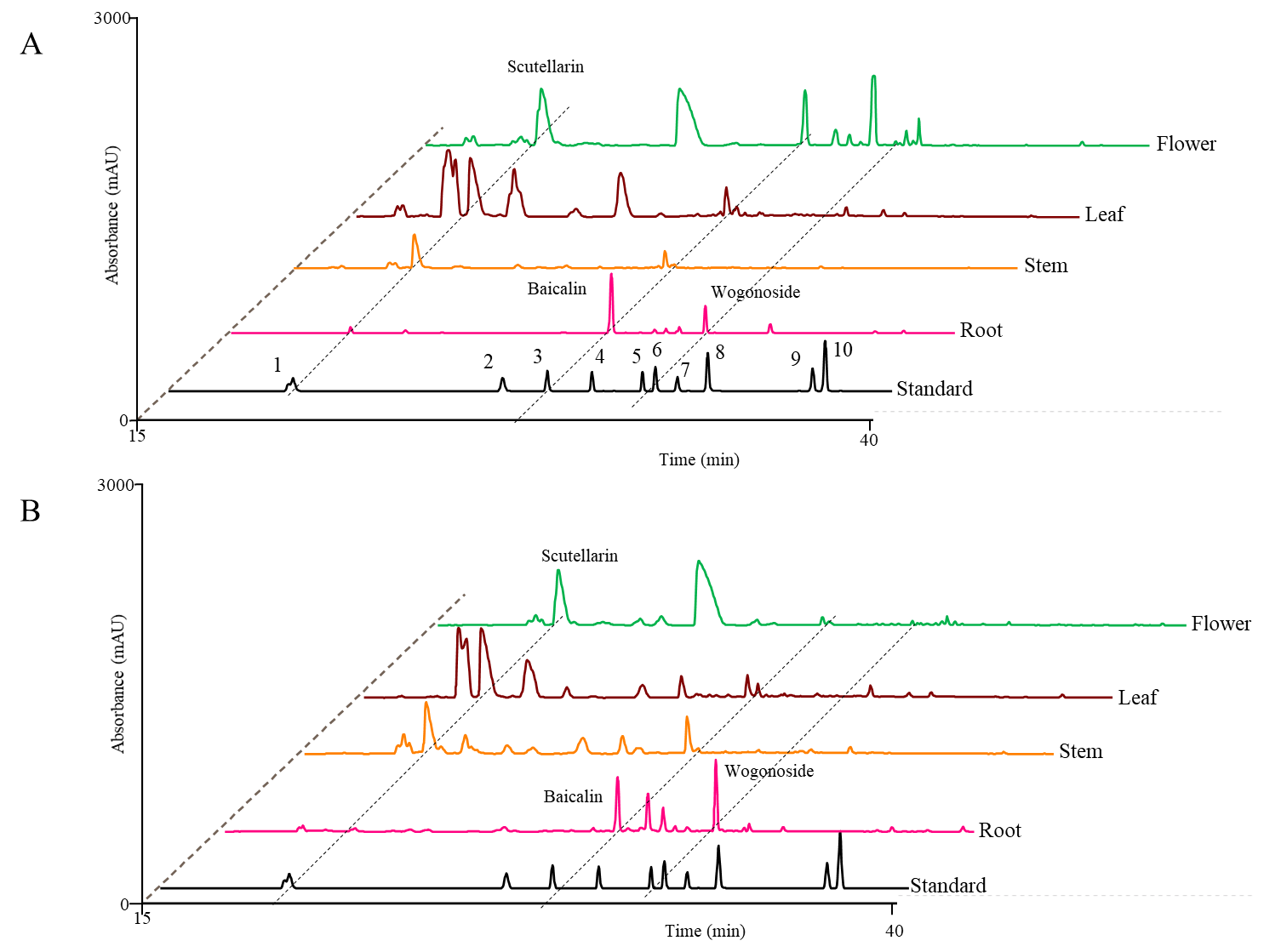
 Supplementary Figure S11. Ultraperformance liduid chromatography (UPLC) detection (280 nm) of flavonoid contents in different tissues of *S. baicalensis* and *S. barbata*, including baicalein, scutellarein, wogonin, and their glycosides (baicalin, scutellarin, and wogonoside). The compound information, including detailed retention times and spectrum data, is listed in Table S11. A. Flavonoid contents of *S. baicalensis*. B. Flavonoid contents of *S. barbata*.
