## Supplemental Figure 12 for "Comparative Genome Analysis of *Scutellaria baicalensis* and *Scutellaria barbata* Reveals the Evolution of Active Flavonoid Biosynthesis"

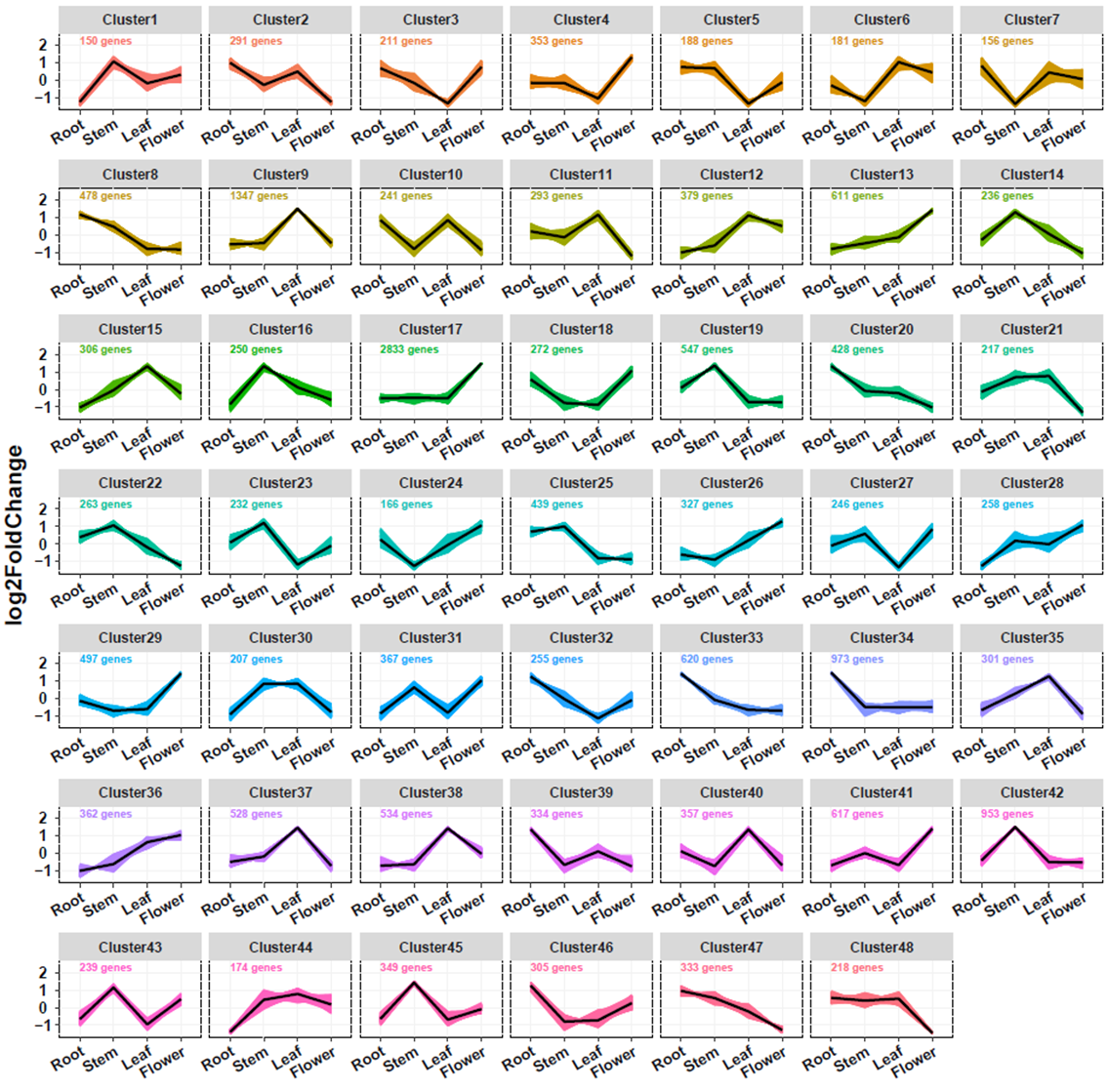

Supplementary Figure S12. All expressed genes were clustered into 48 clusters in different *S. baicalensis* tissues, namely, root, stem, leaf, and flower tissues, based on *k*-means clustering.
