## Supplemental Figure 13 for "Comparative Genome Analysis of *Scutellaria baicalensis* and *Scutellaria barbata* Reveals the Evolution of Active Flavonoid Biosynthesis"

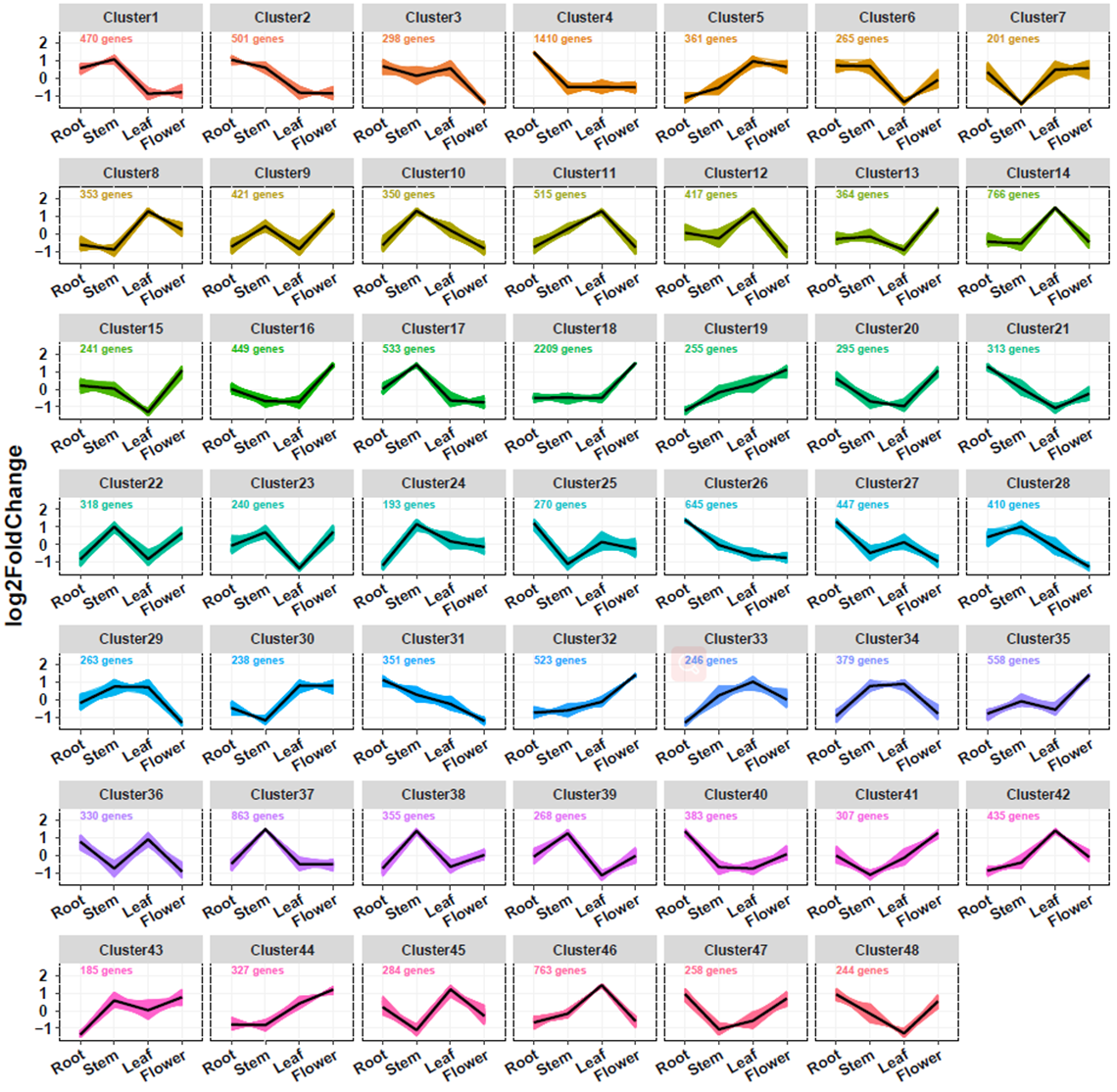

Supplementary Figure S13. All expressed genes were clustered into 48 clusters in different *S. barbata* tissues, namely, root, stem, leaf, and flower tissues, based on k-means clustering.
