## Supplemental Figure 14 for "Comparative Genome Analysis of *Scutellaria baicalensis* and *Scutellaria barbata* Reveals the Evolution of Active Flavonoid Biosynthesis"

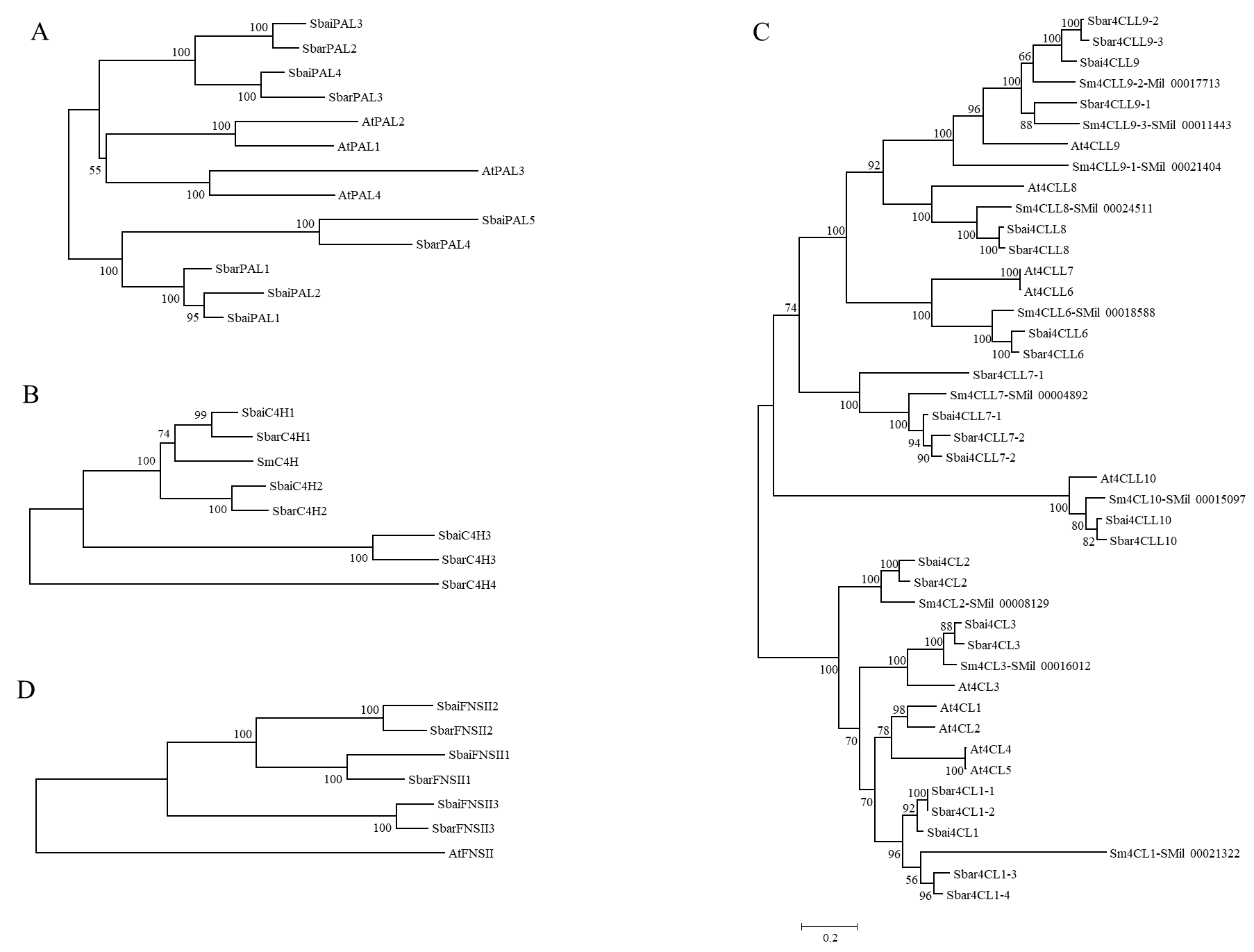

Supplementary Figure S14. Phylogenetic analysis of PAL, C4H, 4CL, and FNSII from *S. baicalensis* and *S. barbata* using the maximum likelihood method.
