## Supplemental Figure 15 for "Comparative Genome Analysis of *Scutellaria baicalensis* and *Scutellaria barbata* Reveals the Evolution of Active Flavonoid Biosynthesis"

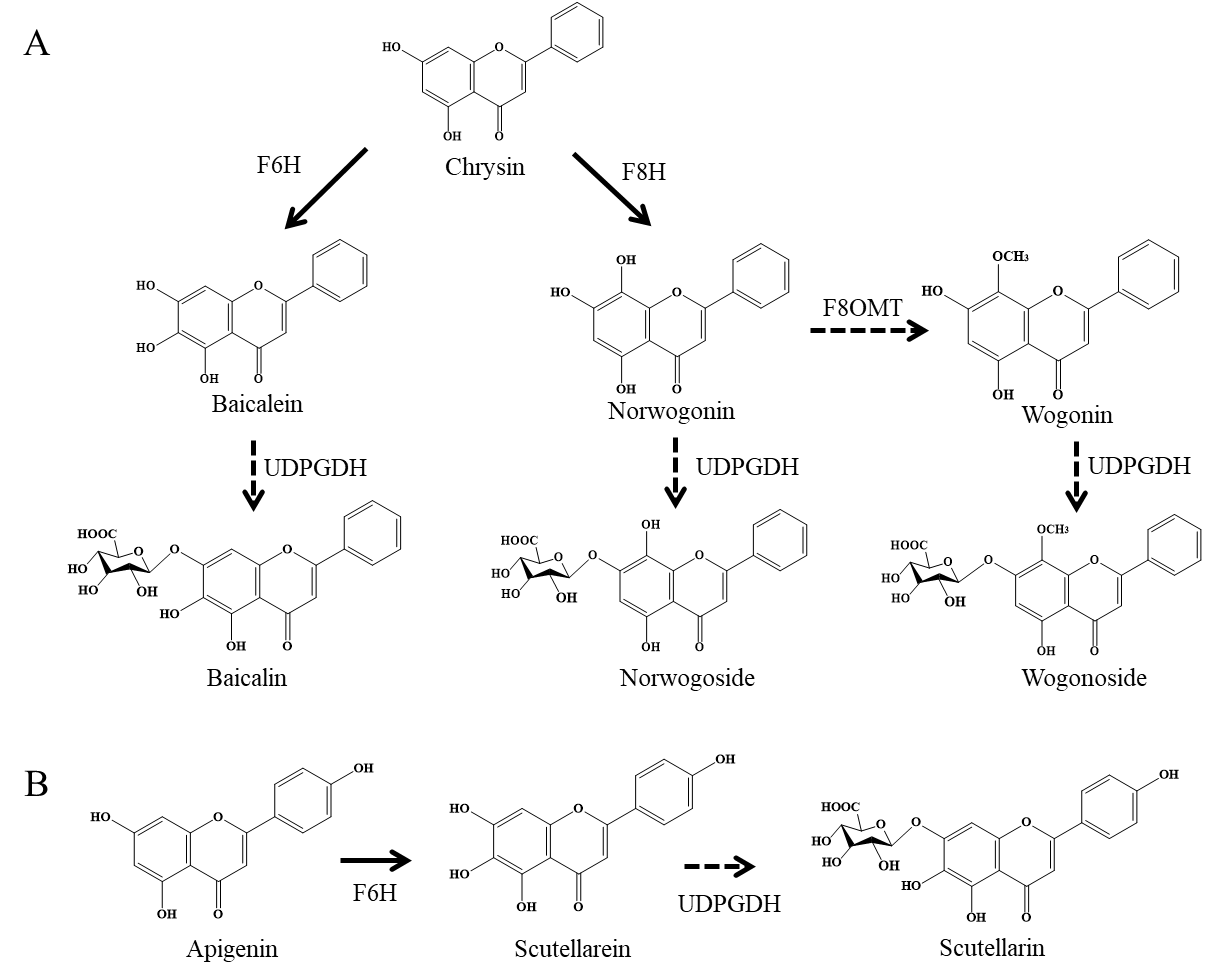

Supplementary Figure S15. The potential biosynthetic pathway of baicalein, scutellarein, wogonin, and their glycosides (baicalin, scutellarin, and wogonoside), catalyzing chrysin and apigenin.
