## Supplementary figures and images for "Comparative Genome Analysis of *Scutellaria baicalensis* and *Scutellaria barbata* Reveals the Evolution of Active Flavonoid Biosynthesis"

### Supplemental Figure 16

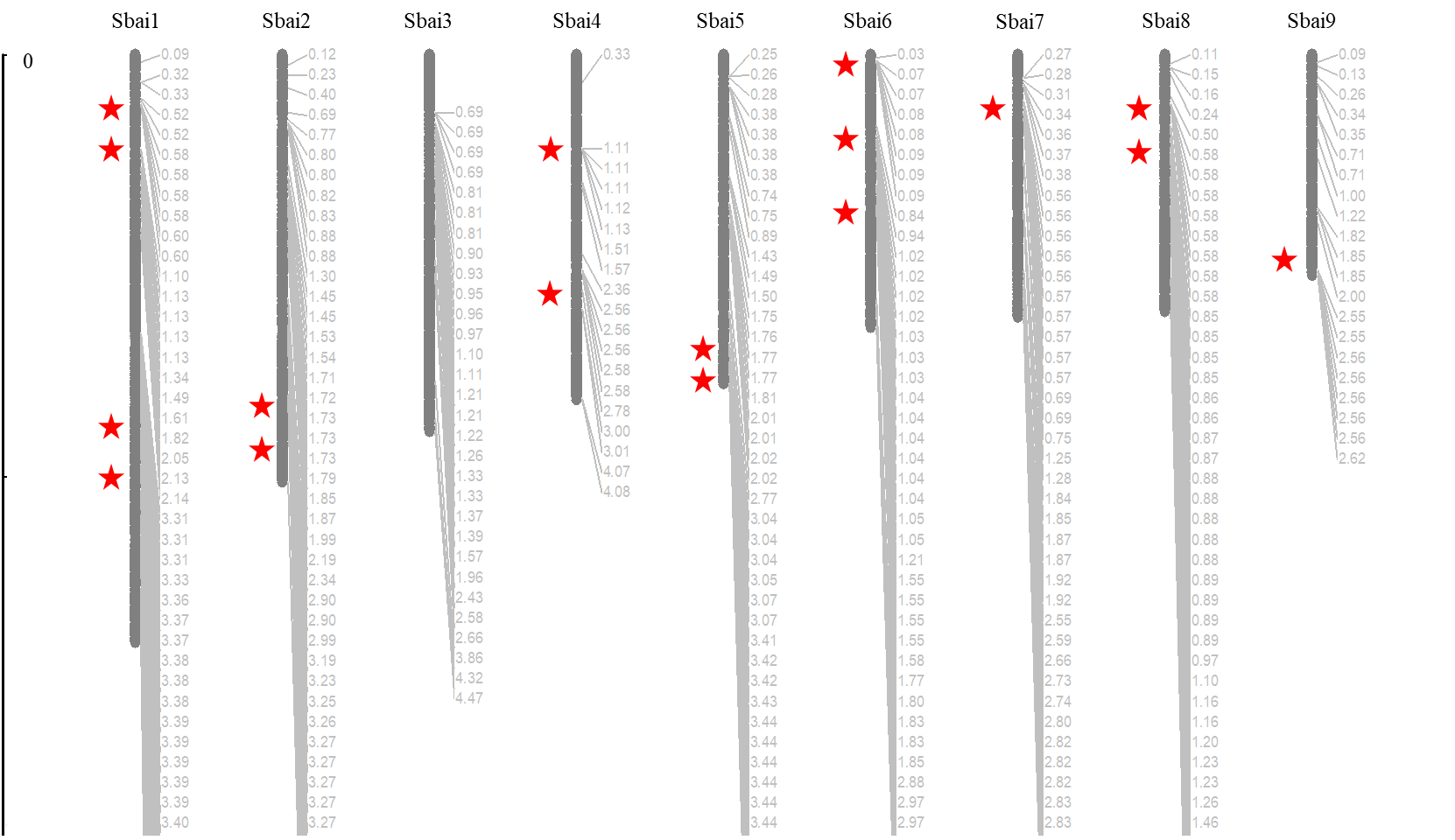

Supplementary Figure S16. The physical clusters of CYP450s (5 gene clusters per 500 kb) in *S. baicalensis*.

### Supplemental Figure 17

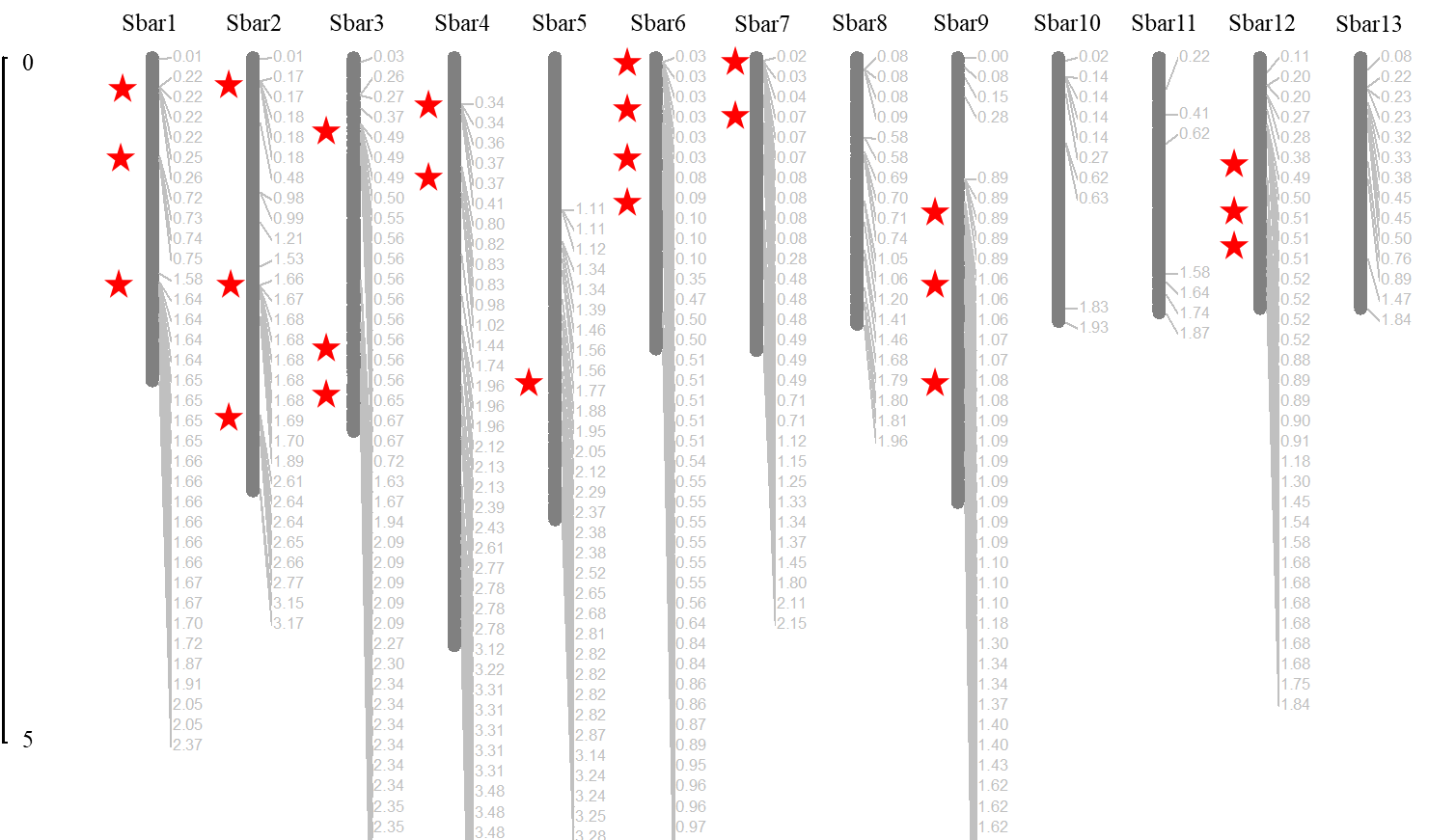

Supplementary Figure S17. The physical clusters of CYP450s (5 gene clusters per 500 kb) in *S. barbata*.
