## Supplemental Figure 18 for "Comparative Genome Analysis of *Scutellaria baicalensis* and *Scutellaria barbata* Reveals the Evolution of Active Flavonoid Biosynthesis"

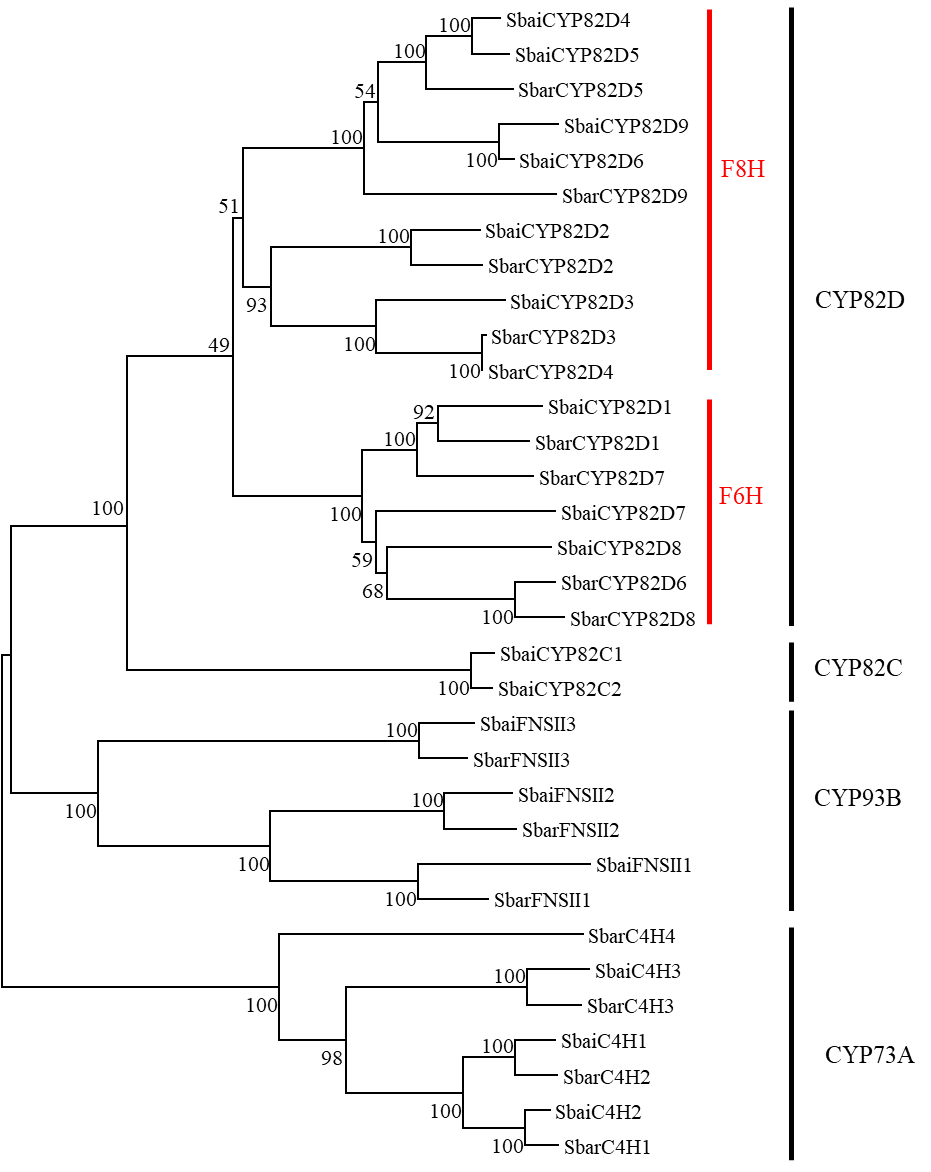

Supplementary Figure S18. The phylogenetic analysis of CYP82D, CYP93B, and CYP73A members from *S. baicalensis* and *S. barbata* using the maximum likelihood method.
