## Supplemental Table 1 for "Comparative Genome Analysis of *Scutellaria baicalensis* and *Scutellaria barbata* Reveals the Evolution of Active Flavonoid Biosynthesis"

Supplementary Table S1. The statistics of sequencing data from the SMRT and ONT platforms and corrected reads using CANU.

|  | *S. baicalensis* (ONT) | *S. barbata* (SMRT) |
| --- | --- | --- |
| Raw reads | 3,193,420 | 8,423,154 |
| Raw data (bp) | 52,035,200,672 | 51,672,515,843 |
| Reads N50 (bp) | 16,324 | 9,843 |
| Filtered reads | 2,761,758 | 7,811,959 |
| Filtered data(bp) | 47,594,396,058 | 483,96,743,827 |
| Reads N50 (bp) | 23,750 | 10,206 |
| Corrected reads using CANU | 567,210 | 1,154,193 |
| Corrected data (bp) | 20,220,782,775 | 18,047,628,482 |
| Reads N50 (bp) | 35,491 | 15,286 |
