## Supplemental Table 2 for "Comparative Genome Analysis of *Scutellaria baicalensis* and *Scutellaria barbata* Reveals the Evolution of Active Flavonoid Biosynthesis"

Supplementary Table S2. The assembled statistics of the *S. baicalensis* and *S. barbata* genome.

|  | *S. baicalensis* | *S. barbata* |
| --- | --- | --- |
| 17-kmer (bp) | 441,861,814 | 404,561,255 |
| Assembled genome(bp) | 376,971,275 | 352,950,734 |
| Assemble genome / predicted genome | 85.3% | 87.2% |
| Contig N50 (bp) | 2,102,880 | 2,496,969 |
| Chromosome | 9 | 13 |
| Chromosomal genome (bp) | 376,437,573 | 348,847,092 |
| Scaffold N50 (bp) | 40,790,749 | 23,709,848 |
| DNA mapping (Illumina reads) | 88.72% | 86.13% |
| RNA mapping (Illumina reads) | 82.97% | 86.56% |
| BUSCO | C:91.5%,F:2.6%,M:5.9%,n:1440 | C:93.0%,F:2.0%,M:5.0%,n:1440 |
