## Supplemental Table 3 for "Comparative Genome Analysis of *Scutellaria baicalensis* and *Scutellaria barbata* Reveals the Evolution of Active Flavonoid Biosynthesis"

Supplementary Table S3. The genome synteny between *S. baicalensis* and *S. barbata*.

| Chromosome | Start | End | Chromosome | Start | End |
| --- | --- | --- | --- | --- | --- |
| Sbai1 | 14,111 | 69,548,543 | Sbar3 | 27,336 | 27,342,083 |
| Sbai1 | 3,220 | 69,557,151 | Sbar7 | 9,799 | 21,455,932 |
| Sbai1 | 15020 | 69566827 | Sbar10 | 23,690 | 19,331,829 |
| Sbai2 | 15,209,838 | 50,693,295 | Sbar4 | 37,696 | 42,993,077 |
| Sbai3 | 48,028 | 44,627,804 | Sbar1 | 356 | 23,705,280 |
| Sbai3 | 48,028 | 44,627,201 | Sbar13 | 23,573 | 18,341,574 |
| Sbai3 | 30,221 | 44,626,995 | Sbar12 | 33,026 | 18,403,481 |
| Sbai3 | 23,017 | 44,627,858 | Sbar11 | 16,453 | 18,713,593 |
| Sbai4 | 70,247 | 10,197,875 | Sbar4 | 2,763 | 43,004,322 |
| Sbai4 | 15,296,756 | 35,686,135 | Sbar4 | 15,318 | 43,001,556 |
| Sbai5 | 31,700 | 38,947,965 | Sbar4 | 18,226 | 43,001,564 |
| Sbai6 | 26,499 | 32,271,471 | Sbar6 | 26,140 | 21,308,016 |
| Sbai7 | 3,811 | 31,044,136 | Sbar2 | 50,952 | 31,645,286 |
| Sbai7 | 93,238 | 31,018,772 | Sbar12 | 24,130 | 18,401,584 |
| Sbai8 | 21,697 | 30,419,592 | Sbar5 | 18 | 33,852,674 |
| Sbai8 | 23,848 | 30,421,994 | Sbar9 | 1 | 37,631,259 |
| Sbai9 | 40,155 | 26,199,817 | Sbar8 | 38,466 | 19,541,103 |
| Sbai9 | 35,238 | 20,977,779 | Sbar6 | 7,066 | 21,308,013 |
