## Supplemental Table 4 for "Comparative Genome Analysis of *Scutellaria baicalensis* and *Scutellaria barbata* Reveals the Evolution of Active Flavonoid Biosynthesis"

Supplementary Table S4. Genome annotations among *S. baicalensis*, *S. barbata* and *S. miltiorrhiza*.

| **Annotation** | ***S. baicalensis*** | ***S. barbata*** | ***S. miltiorrhiza*** |
| --- | --- | --- | --- |
| No. of predicted transcripts and proteins | 33,414 | 41,697 | 30,478 |
| Average gene length (bp) | 4,553 | 2,257 | 2,826 |
| Average CDS length (bp) | 1,130 | 1,137 | 1,173 |
| Exon Number | 161,060 | 182,344 | 164,031 |
| Average exon length (bp) | 234 | 260 | 228 |
| Average intron length (bp) | 296 | 337 | 268 |
| GC content of transcripts (%) | 44.92 | 45.34 | 47.96 |
| Percentage of whole gene length in genome (%) | 10.01 | 13.43 | 6.65 |
| BUSCO | C:87.3%,F:5.9%,M:6.8% | C:89.7%,F:4.6%,M:5.7% | C:84.6%,F:6.5%,M:8.9%,n:1375 |
| Masked repeat sequence length (bp) | 208,004,279 | 188,790,851 | 292,797,272 |
| Percentage of repeat sequences in genome (%) | 55.17 | 53.49 | 54.44 |
