## Supplemental Table 5 for "Comparative Genome Analysis of *Scutellaria baicalensis* and *Scutellaria barbata* Reveals the Evolution of Active Flavonoid Biosynthesis"

Supplementary Table S5. Annotation of *S. baicalensis* TEs.

| Repeat Class | Elements number | Length occupied (bp) | Percentage of sequence |
| --- | --- | --- | --- |
| **Retrotransposon** | **413,071** | **202,569,821** | **53.73 %** |
| **RNA transposable elements** | **87,768** | **82,875,127** | **32.73 %** |
| non-LTR(Long terminal repeat) | 9,787 | 3,547,610 | 0.94 % |
| LINE | 7,183 | 3,023,969 | 0.80 % |
| SINE | 2,604 | 52,3641 | 0.14 % |
| LTR | 179,033 | 119,831,312 | 31.79% |
| Gypsy | 36,704 |  |  |
| Copia | 43,124 |  |  |
| **DNA elements** | **90,835** | **39,583,481** | **10.49%** |
| **Unclassified TEs** | **133,416** | **39,607,418** | **10.50%** |
| Small RNA | 292 | 216,778 | 0.06 % |
| Satellites | 1,347 | 674,788 | 0.18 % |
| Simple repeats | 106,867 | 4,836,933 | 1.28 % |
| Low complexity | 16,133 | 830,806 | 0.22 % |
| **Total Repeats** |  | **208,004,279** | 55.17 % |
