## Supplemental Table 6 for "Comparative Genome Analysis of *Scutellaria baicalensis* and *Scutellaria barbata* Reveals the Evolution of Active Flavonoid Biosynthesis"

Supplementary Table S6. Annotation of *S. barbata* TEs.

| Repeat Class | Elements number | Length occupied (bp) | Percentage of sequence |
| --- | --- | --- | --- |
| **Retrotransposon** | **304,091** | **182,957,926** | **51.83 %** |
| **RNA transposable elements** | **72,633** | **98,481,941** | **32.75%** |
| non-LTR(Long terminal repeat) | 4,969 | 2,462,125 | 0.70 % |
| LINE | 4,080 | 2,157,913 | 0.61 % |
| SINE | 889 | 304,212 | 0.09 % |
| LTR | 105,267 | 113,115,881 | 32.05% |
| Gypsy | 36,745 |  |  |
| Copia | 35,064 |  |  |
| **DNA elements** | **66,220** | **30,502,000** | **8.64 %** |
| **Unclassified TEs** | **127,635** | **36,877,920** | **10.44%** |
| Small RNA | 155 | 65,333 | 0.02 % |
| Satellites | 799 | 491,615 | 0.14 % |
| Simple repeats | 110,331 | 4,993,703 | 1.41 % |
| Low complexity | 18,197 | 894,549 | 0.25 % |
| **Total Repeats** |  | **188,790,851** | 53.49 % |
