## Supplemental Table 7 for "Comparative Genome Analysis of *Scutellaria baicalensis* and *Scutellaria barbata* Reveals the Evolution of Active Flavonoid Biosynthesis"

Supplementary Table S7. Summary of intact LTR retrotransposons in *S. baicalensis* and *S. barbata*.

|  | *S. baicalensis* | *S. barbata* |
| --- | --- | --- |
| Intact LTRs | 1225 | 1654 |
| Average inserting times of all intact LTRs (MYA) | 1.41 | 0.88 |
| *Gypsy* | 342 | 310 |
| Max insertion times of *Gypsy* (MYA) | 4.14 | 4.39 |
| Min insertion times of *Gypsy* (MYA) | 0.072 | 0.015 |
| Average insertion times of *Gypsy* (MYA) | 1.42 | 0.96 |
| *Copia* | 354 | 618 |
| Max insertion times of *Copia* (MYA) | 6.92 | 3.87 |
| Min insertion times of *Copia* (MYA) | 0.14 | 0.047 |
| Average insertion times of *Copia* (MYA) | 1.55 | 0.90 |
