## Supplemental Table 8 for "Comparative Genome Analysis of *Scutellaria baicalensis* and *Scutellaria barbata* Reveals the Evolution of Active Flavonoid Biosynthesis"

Supplementary Table S8. Annotation of *S. baicalensis* and *S. barbata* rRNA.

|  | *S. baicalensis* | *S. barbata* |
| --- | --- | --- |
| rRNA | 813 | 210 |
| 8s rRNA | 785 | 189 |
| 18s rRNA | 16 | 8 |
| 28s rRNA | 12 | 13 |
