## Supplemental Table 9 for "Comparative Genome Analysis of *Scutellaria baicalensis* and *Scutellaria barbata* Reveals the Evolution of Active Flavonoid Biosynthesis"

Supplementary Table S9. Identification of SSRs in the *S. baicalensis* and *S. barbata* genome.

| Types | Unit size | Cut-off | Number of SSR in *S. baicalensis* | Number of SSR in *S. barbata* |
| --- | --- | --- | --- | --- |
| Monomer | 1 | 10 | 73,270 | 93,162 |
| Dimer | 2 | 6 | 58,417 | 42,280 |
| Trimer | 3 | 5 | 9,876 | 10,742 |
| Tetramer | 4 | 5 | 740 | 1,160 |
| Pentamer | 5 | 5 | 373 | 213 |
| Hexamer | 6 | 5 | 275 | 148 |
| Total |  |  | 142,951 | 147,705 |
