## Supplemental Table 10 for "Comparative Genome Analysis of *Scutellaria baicalensis* and *Scutellaria barbata* Reveals the Evolution of Active Flavonoid Biosynthesis"

Supplementary Table S10. Gene family expansion and contraction of candidate species according to phylogenetic analysis (*P* < 0.01).

| Phylogenetic tree node | Expansion | Remain | Contraction |
| --- | --- | --- | --- |
| 0 | 2176 | 16237 | 964 |
| 2 | 965 | 15169 | 3344 |
| 1 | 1639 | 10281 | 7552 |
| 13 | 475 | 16475 | 2527 |
| 4 | 1180 | 16697 | 1599 |
| 6 | 1853 | 15993 | 1632 |
| 5 | 693 | 17055 | 1727 |
| 8 | 2543 | 12664 | 4271 |
| 7 | 109 | 18813 | 550 |
| 10 | 5117 | 11055 | 3305 |
| 9 | 1145 | 16699 | 1629 |
| 12 | 6388 | 9067 | 4023 |
| 11 | 856 | 17742 | 878 |
| 17 | 47 | 16797 | 2634 |
| 14 | 2142 | 13014 | 4320 |
| 16 | 5934 | 11843 | 1699 |
| 15 | 43 | 19417 | 11 |
| 18 | 1306 | 13251 | 4921 |
| 3 | 143 | 19125 | 207 |
| 20 | 1104 | 7030 | 11344 |
