## Supplemental Table 11 for "Comparative Genome Analysis of *Scutellaria baicalensis* and *Scutellaria barbata* Reveals the Evolution of Active Flavonoid Biosynthesis"

Supplementary Table S11. The *Ks* value and divergence time of paralogous or orthologous gene pairs among *S. baicalensis*, *S. barbata*, *S. miltiorrhiza*, *S. indicum*, and *V. vinifera*.

| Species | Species | Orthologous / Paralogous | *Ks* peak | Divergence time for speciation (MYA) | WGD time (MYA) | Synonymous substitutions per site per MYA |
| --- | --- | --- | --- | --- | --- | --- |
| *S. baicalensis* | *S. baicalensis* | 5978 | 0.886467515 | N/A | 60.71031093 | N/A |
| *S. barbata* | *S. barbata* | 5366 | 0.862984932 | N/A | 59.10209078 | N/A |
| *S. miltiorrhiza* | *S. miltiorrhiza* | 5764 | 1.021492368 | N/A | 69.95757681 | N/A |
| *S. indicum* | *S. indicum* | 6702 | 0.675124266 | N/A | 46.23632949 | N/A |
| *V. vinifera* | *V. vinifera* | 1993 | 1.203482387 | N/A | N/A | N/A |
| *S. baicalensis* | *S. miltiorrhiza* | 11028 | 0.598805871 | 41.009614 | N/A | 0.007300799 |
| *S. baicalensis* | *S. indicum* | 11712 | 0.604676517 | 49.897716 | N/A | 0.006059161 |
| *S. baicalensis* | *V. vinifera* | 6098 | 1.526367906 | 115.832779 | N/A | 0.00658867 |
| *S. barbata* | *S. miltiorrhiza* | 11123 | 0.581193933 | 41.009614 | N/A | 0.00708607 |
| *S. barbata* | *S. indicum* | 11936 | 0.557711350 | 49.897716 | N/A | 0.005588546 |
| *S. barbata* | *V. vinifera* | 6282 | 1.514626614 | 115.832779 | N/A | 0.006537988 |
| *S. baicalensis* | *S. barbata* | 16204 | 0.16437802 | 13.281676 | N/A | 0.006188151 |
