## Supplemental Table 12 for "Comparative Genome Analysis of *Scutellaria baicalensis* and *Scutellaria barbata* Reveals the Evolution of Active Flavonoid Biosynthesis"

Supplementary Table S12. The compound information of UPLC detection including retention time and spectrum, is shown in Figure S8.

|  | Compounds | Time (min) | λmax (nm) | MW |
| --- | --- | --- | --- | --- |
| 1 | Scutellarin | 19.303 | 282.50 | 462.37 |
| 2 | Scutellarein | 26.553 | 283.20 | 286.24 |
| 3 | Baicalin | 28.097 | 277.50 | 446.37 |
| 4 | Norwogoside | 29.637 | 279.97 | 446.36 |
| 5 | Wogonoside | 31.383 | 274.32 | 460.39 |
| 6 | Apigenin | 31.827 | 267.26 | 270.24 |
| 7 | Norwogonin | 32.590 | 280.41 | 270.24 |
| 8 | Baicalein | 33.637 | 275.72 | 270.24 |
| 9 | Wogonin | 37.263 | 275.23 | 284.27 |
| 10 | Chrysin | 37.697 | 265.65 | 254.24 |
