## Supplemental Table 15 for "Comparative Genome Analysis of *Scutellaria baicalensis* and *Scutellaria barbata* Reveals the Evolution of Active Flavonoid Biosynthesis"

Supplementary Table S15. The expression of chrysin and apigenin biosynthetic genes in different organs of *S. baicalensis*.

| Gene name | Gene ID | Root | Stem | Leaf | Flower |
| --- | --- | --- | --- | --- | --- |
| SbaiPAL1 | Sbai1A386T84 | 20.83 | 50.64 | 0.59 | 1.34 |
| SbaiPAL2 | Sbai1A383T80 | 8.43 | 25.79 | 0.16 | 0.30 |
| SbaiPAL3 | Sbai4A267T179 | 468.51 | 921.80 | 330.78 | 192.69 |
| SbaiPAL4 | Sbai3A247T84 | 45.46 | 57.20 | 90.06 | 6.42 |
| SbaiPAL5 | Sbai1A614T103 | 10.04 | 4.31 | 0.18 | 0.02 |
| Sbai4CL1 | Sbai5A101T78 | 103.00 | 176.58 | 9.10 | 5.57 |
| Sbai4CL2 | Sbai2A267T110 | 1.24 | 1.06 | 1.63 | 16.03 |
| Sbai4CL3 | Sbai1A162T99 | 42.26 | 47.74 | 120.93 | 23.66 |
| Sbai4CLL6 | Sbai8A100T152 | 24.47 | 1.38 | 87.51 | 36.67 |
| Sbai4CLL7-1 | Sbai7C308T10 | 29.35 | 14.63 | 26.64 | 15.29 |
| Sbai4CLL7-2 | Sbai7C308T11 | 0.00 | 0.00 | 0.00 | 2.50 |
| Sbai4CLL8 | Sbai6A97T88 | 213.72 | 330.55 | 352.63 | 14.09 |
| Sbai4CLL9 | Sbai2A90T96 | 0.10 | 7.86 | 33.75 | 1.65 |
| Sbai4CLL10 | Sbai2A87T48 | 275.12 | 85.67 | 155.36 | 72.28 |
| SbaiCHS1 | Sbai7C107T21 | 162.34 | 2027.01 | 2395.79 | 222.30 |
| SbaiCHS2 | Sbai7C102T21 | 35.04 | 5.39 | 10.79 | 1.84 |
| SbaiCHS3 | Sbai7C100T5 | 29.68 | 8.81 | 7.56 | 2.28 |
| SbaiCHS4 | Sbai7A118T54 | 74.65 | 54.77 | 29.89 | 25.79 |
| SbaiCHS5 | Sbai7C104T3 | 35.43 | 9.74 | 8.94 | 2.82 |
| SbaiCHS6 | Sbai3C104T3 | 0.00 | 0.94 | 12.47 | 0.05 |
| SbaiCHS7 | Sbai8A22T136 | 0.00 | 0.00 | 0.05 | 0.09 |
| SbaiCHS8 | Sbai8C66T3 | 0.02 | 1.05 | 0.43 | 1.08 |
| SbaiCHI | Sbai3C171T4 | 159.46 | 129.39 | 100.72 | 358.75 |
| SbaiFNSII1 | Sbai3P121T26 | 8.71 | 9.66 | 36.53 | 55.25 |
| SbaiFNSII2 | Sbai3A121T93 | 92.37 | 121.61 | 2.78 | 13.52 |
| SbaiFNSII3 | Sbai4A257T106 | 0.79 | 0.00 | 0.03 | 0.00 |
| C4H1 | Sbai3A132T111 | 66.80 | 374.44 | 442.17 | 130.62 |
| C4H2 | Sbai8C150T8 | 8.71 | 9.66 | 36.53 | 30.45 |
| C4H3 | Sbai8A150T117 | 6.95 | 3.08 | 3.91 | 15.65 |
