## Supplemental Table 16 for "Comparative Genome Analysis of *Scutellaria baicalensis* and *Scutellaria barbata* Reveals the Evolution of Active Flavonoid Biosynthesis"

Supplementary Table S16. The expression of chrysin and apigenin biosynthetic genes in different organs of *S. barbata*.

| Gene name | Gene ID | Root | Stem | Leaf | Flower |
| --- | --- | --- | --- | --- | --- |
| SbarPAL1 | Sbar5A218T212 | 18.55 | 68.76 | 16.08 | 62.55 |
| SbarPAL2 | Sbar4A4T132 | 147.95 | 418.11 | 251.03 | 222.89 |
| SbarPAL3 | Sbar11A170T161 | 83.38 | 84.76 | 61.14 | 19.06 |
| SbarPAL4 | Sbar10A150T96 | 6.81 | 2.81 | 0.78 | 0.11 |
| Sbar4CL1-1 | Sbar4C297T22 | 1.16 | 1.78 | 0.98 | 0.53 |
| Sbar4CL1-2 | Sbar4C298T24 | 7.10 | 8.72 | 2.64 | 2.11 |
| Sbar4CL1-3 | Sbar3A77T120 | 0.63 | 0.59 | 0.01 | 0.01 |
| Sbar4CL1-4 | Sbar9A326T113 | 25.62 | 5.18 | 0.85 | 1.68 |
| Sbar4CL2 | Sbar5A256T179 | 67.84 | 29.78 | 22.22 | 159.44 |
| Sbar4CL3 | Sbar4A380T150 | 223.87 | 243.07 | 186.02 | 62.82 |
| Sbar4CLL6 | Sbar9A255T177 | 0.26 | 5.40 | 63.86 | 10.23 |
| Sbar4CLL7-1 | Sbar13A3T207 | 32.03 | 23.25 | 36.33 | 35.24 |
| Sbar4CLL7-2 | Sbar3A216T75 | 0.00 | 0.30 | 0.29 | 3.34 |
| Sbar4CLL8 | Sbar6A181T46 | 40.42 | 143.13 | 275.04 | 14.78 |
| Sbar4CLL9-1 | Sbar4A312T146 | 0.37 | 0.24 | 0.29 | 3.03 |
| Sbar4CLL9-2 | Sbar9C331T22 | 1.76 | 0.49 | 13.25 | 10.37 |
| Sbar4CLL9-3 | Sbar9A331T186 | 0.02 | 5.62 | 175.03 | 3.84 |
| Sbar4CLL10 | Sbar5A154T110 | 602.54 | 392.44 | 517.89 | 49.31 |
| SbarCHS1 | Sbar2C282T9 | 3098.44 | 5287.19 | 3673.26 | 1562.77 |
| SbarCHS2 | Sbar5A272T111 | 0.84 | 3.30 | 0.50 | 3.20 |
| SbarCHS3 | Sbar5A316T184 | 0.12 | 0.77 | 0.23 | 34.47 |
| SbarCHI | Sbar1A223T166 | 511.65 | 538.07 | 315.10 | 330.28 |
| SbarFNSII1 | Sbar13C32T13 | 10.55 | 87.50 | 78.56 | 340.41 |
| SbarFNSII2 | Sbar13A32T113 | 215.62 | 28.64 | 0.65 | 0.29 |
| SbarFNSII3 | Sbar4C196T19 | 9.31 | 0.48 | 0.13 | 0.19 |
| SbarC4H1 | Sbar9A209T130 | 110.87 | 389.91 | 195.03 | 168.72 |
| SbarC4H2 | Sbar13C44T18 | 126.39 | 96.22 | 21.43 | 150.01 |
| SbarC4H3 | Sbar9A209T128 | 0.30 | 0.91 | 2.53 | 0.15 |
| SbarC4H4 | Sbar4A98T57 | 1.17 | 0.37 | 0.00 | 0.00 |
