## Supplemental Table 17 for "Comparative Genome Analysis of *Scutellaria baicalensis* and *Scutellaria barbata* Reveals the Evolution of Active Flavonoid Biosynthesis"

Supplementary Table S17. The *Ka* and *Ks* analysis of chrysin and apigenin biosynthetic genes in *S. baicalensis* and *S. barbata*.

| Enzymes | *S. baicalensis* | *S. barbata* | Ka | Ks | Ka/Ks |
| --- | --- | --- | --- | --- | --- |
| C4H | Sbai3A132T111 | Sbar13C44T18 | 0.0166787 | 0.21811 | 0.0764692 |
|  | Sbai8A150T117 | Sbar4A98T57 | 0.0362953 | 0.29481 | 0.123114 |
| PAL | Sbai1A386T84 | Sbar5A218T212 | 0.0103182 | 0.276419 | 0.0373279 |
|  | Sbai4A267T179 | Sbar4A4T132 | 0.0124043 | 0.303438 | 0.0408792 |
|  | Sbai3A247T84 | Sbar11A170T161 | 0.0182678 | 0.51455 | 0.0355025 |
|  | Sbai1A614T103 | Sbar10A150T96 | 0.0583147 | 0.246815 | 0.236268 |
| 4CL | Sbai2A267T110 | Sbar5A256T179 | 0.0501394 | 0.284728 | 0.176096 |
|  | Sbai1A162T99 | Sbar4A380T150 | 0.0333036 | 0.315783 | 0.105464 |
|  | Sbai8A100T152 | Sbar9A255T177 | 0.0345797 | 0.15462 | 0.223643 |
|  | Sbai2A90T96 | Sbar9A331T186 | 0.101077 | 0.250834 | 0.402965 |
|  | Sbai2A87T48 | Sbar5A154T110 | 0.0199619 | 0.182408 | 0.109436 |
|  | Sbai7C308T11 | Sbar13A3T207 | 0.0497812 | 0.176752 | 0.281645 |
| CHS | Sbai7C107T21 | Sbar2C282T9 | 0.0380835 | 1.18405 | 0.0321638 |
|  | Sbai8A22T136 | Sbar5A316T184 | 0.0303051 | 0.278844 | 0.108681 |
|  | Sbai8C66T3 | Sbar5A272T111 | 0.027692 | 0.41085 | 0.0674017 |
| CHI | Sbai3C171T4 | Sbar1A223T166 | 0.0365224 | 0.29818 | 0.122485 |
| FNSII | Sbai3A121T93 | Sbar13A32T113 | 0.0467232 | 0.805128 | 0.058032 |
|  | Sbai4A257T106 | Sbar4C196T19 | 0.0377574 | 0.165362 | 0.228333 |
