## Supplemental Table 18 for "Comparative Genome Analysis of *Scutellaria baicalensis* and *Scutellaria barbata* Reveals the Evolution of Active Flavonoid Biosynthesis"

Supplementary Table S18. The expression of CYP82D members in different tissues of *S. baicalensis* and *S. barbata*.

|  | Gene name | Gene ID | Root | Stem | Leaf | Flower |
| --- | --- | --- | --- | --- | --- | --- |
| *S. baicalensis* | SbaiCYP82D1 | Sbai6A9T110 | 29.92 | 3.18 | 0.68 | 69.64 |
|  | SbaiCYP82D2 | Sbai1A57T82 | 36.79 | 0.82 | 8.89 | 0.12 |
|  | SbaiCYP82D3 | Sbai1C57T21 | 11.21 | 6.81 | 11.18 | 6.61 |
|  | SbaiCYP82D4 | Sbai1A33T134 | 7.23 | 9.36 | 6.77 | 0.12 |
|  | SbaiCYP82D5 | Sbai1A58T73 | 2.28 | 32.91 | 29.33 | 2.59 |
|  | SbaiCYP82D6 | Sbai1C459T13 | 8.36 | 0.00 | 0.00 | 0.00 |
|  | SbaiCYP82D7 | Sbai6A9T108 | 0.00 | 0.00 | 0.00 | 0.00 |
|  | SbaiCYP82D8 | Sbai6C9T2 | 10.93 | 26.00 | 48.90 | 2.72 |
|  | SbaiCYP82D9 | Sbai1A457T74 | 1.43 | 0.00 | 0.00 | 0.00 |
| *S. barbata* | SbarCYP82D1 | Sbar6C10T12.1 | 10.70 | 30.11 | 38.46 | 0.75 |
|  | SbarCYP82D2 | Sbar7A134T46.1 | 5.62 | 5.07 | 2.87 | 0.09 |
|  | SbarCYP82D3 | Sbar7A133T50.1 | 0.22 | 2.42 | 4.62 | 0.04 |
|  | SbarCYP82D4 | Sbar7A137T41.1 | 1.18 | 6.03 | 5.49 | 0.02 |
|  | SbarCYP82D5 | Sbar7A114T56.1 | 85.87 | 14.33 | 1.22 | 2.18 |
|  | SbarCYP82D7 | Sbar3C37T16.1 | 0.00 | 0.22 | 0.00 | 0.89 |
|  | SbarCYP82D6 | Sbar6A10T128.1 | 185.52 | 61.96 | 186.00 | 1.61 |
|  | SbarCYP82D8 | Sbar6A10T127.1 | 447.96 | 103.55 | 623.09 | 11.68 |
|  | SbarCYP82D9 | Sbar7A112T53.1 | 179.86 | 56.66 | 40.44 | 75.29 |
|  | SbarCYP82D10 | Sbar12A130T67.1 | 0.00 | 0.06 | 0.03 | 0.08 |
