## Supplemental Table 19 for "Comparative Genome Analysis of *Scutellaria baicalensis* and *Scutellaria barbata* Reveals the Evolution of Active Flavonoid Biosynthesis"

Supplementary Table S19. The *Ks* values of gene pairs related to flavone biosynthesis in *S. baicalensis* and *S. barbata*.

|  | Paralogous pairs | Ks |
| --- | --- | --- |
| *S. baicalensis* | SbaiPAL1 – SbaiPAL2 | 0.0440141 |
|  | SbaiPAL3 – SbaiPAL4 | 1.82591 |
|  | Sbai4CL2 – Sbai4CL3 | 3.17105 |
|  | Sbai4CLL7-1–Sbai4CLL7-2 | 0.251031 |
|  | SbaiCHS4 – SbaiCHS3 | 0.0806434 |
|  | SbaiCHS3 – SbaiCHS5 | 0.00586652 |
|  | SbaiCHS5 – SbaiCHS4 | 0.0825493 |
|  | SbaiCHS5 – SbaiCHS1 | 0.379022 |
|  | SbaiCHS1 – SbaiCHS4 | 0.368591 |
|  | SbaiCHS7 – SbaiCHS8 | 3.13754 |
|  | SbaiFNS1 – SbaiFNS2 | 1.94371 |
|  | SbaiC4H1 – SbaiC4H2 | 1.69773 |
|  | SbaiCYP82D8 – SaiCYP82D1 | 0.87768 |
|  | SbaiCYP82D8 – SbaiCYP82D7 | 0.634991 |
|  | SbaiCYP82D2 – SbaiCYP82D3 | 0.942118 |
|  | SbaiCYP82D4 – SbaiCYP82D5 | 0.0803808 |
|  | SbaiCYP82D6 – SbaiCYP82D9 | 0.0707554 |
| *S. barbata* | SbarPAL2 – SbarPAL3 | 1.06785 |
|  | Sbar4CL1-1 – Sbar4CL1-2 | 0 |
|  | Sbar4CL1-3 – Sbar4CL1-4 | 0.177134 |
|  | Sbar4CLL9-2 –Sbar4CLL9-3 | 0.0651871 |
|  | SbarCHS2 – SbarCHS3 | 3.07387 |
|  | SbarFNS1 – SbarFNS2 | 1.14678 |
|  | SbarC4H1 – SbarC4H2 | 1.38873 |
|  | SbarCYP82D8 – SbarCYP82D1 | 0.763061 |
|  | SbarCYP82D8 – SbarCYP82D7 | 0.629526 |
|  | SbarCYP82D8 – SbarCYP82D6 | 0.0559024 |
|  | SbarCYP82D2 – SbarCYP82D3 | 1.02687 |
|  | SbarCYP82D3 – SbarCYP82D4 | 0.0115003 |
|  | SbarCYP82D5 – SbarCYP82D9 | 0.335215 |
